# A novel RNA-binding activity of ECD contributes to U5 snRNP stability and pre-mRNA splicing

**DOI:** 10.1101/2025.01.24.634785

**Authors:** Mohsin Raza, Asher Rajkumar Rajan, Achyuth Kalluchi, Bhopal Mohapatra, Irfana Saleem, Benjamin B. Kennedy, Kishor K. Bhakat, Hamid Band, M. Jordan Rowley, Vimla Band

## Abstract

Human ecdysoneless protein (ECD) plays an essential role in regulating cell cycle progression and cell survival. ECD has previously been implicated in RNA splicing through its association with spliceosomal proteins. Here, using EMSA, fluorescence polarization assays, and mutational analysis, we demonstrate that ECD directly binds to RNA. Enhanced CLIP-seq analysis identified a broad repertoire of mRNAs bound to ECD in cells. RNA-seq analyses revealed that ECD depletion leads to widespread splicing aberrations and altered gene expression. ECD binding to RNAs was enriched near splice sites, and a substantial fraction of ECD-bound transcripts exhibited splicing defects upon ECD depletion. ECD associates with and stabilizes the U5 snRNP complex specific proteins. While depletion of ECD reduced the levels of key U5-specifc proteins, these proteins exhibited an increased association with the R2TP complex in knockout cells. Notably, we found ECD to directly bind to U5 snRNA, and an RNA binding defective mutant of ECD (Δ135–148) failed to rescue the reduced levels of U5-specific proteins or the proliferation defect induced by ECD depletion. Collectively, these findings demonstrate that ECD binds to RNAs, including the U5 snRNA, and that RNA-binding is required for ECD to stabilize the U5 snRNP and for cellular functions.

**Graphical Abstract:** 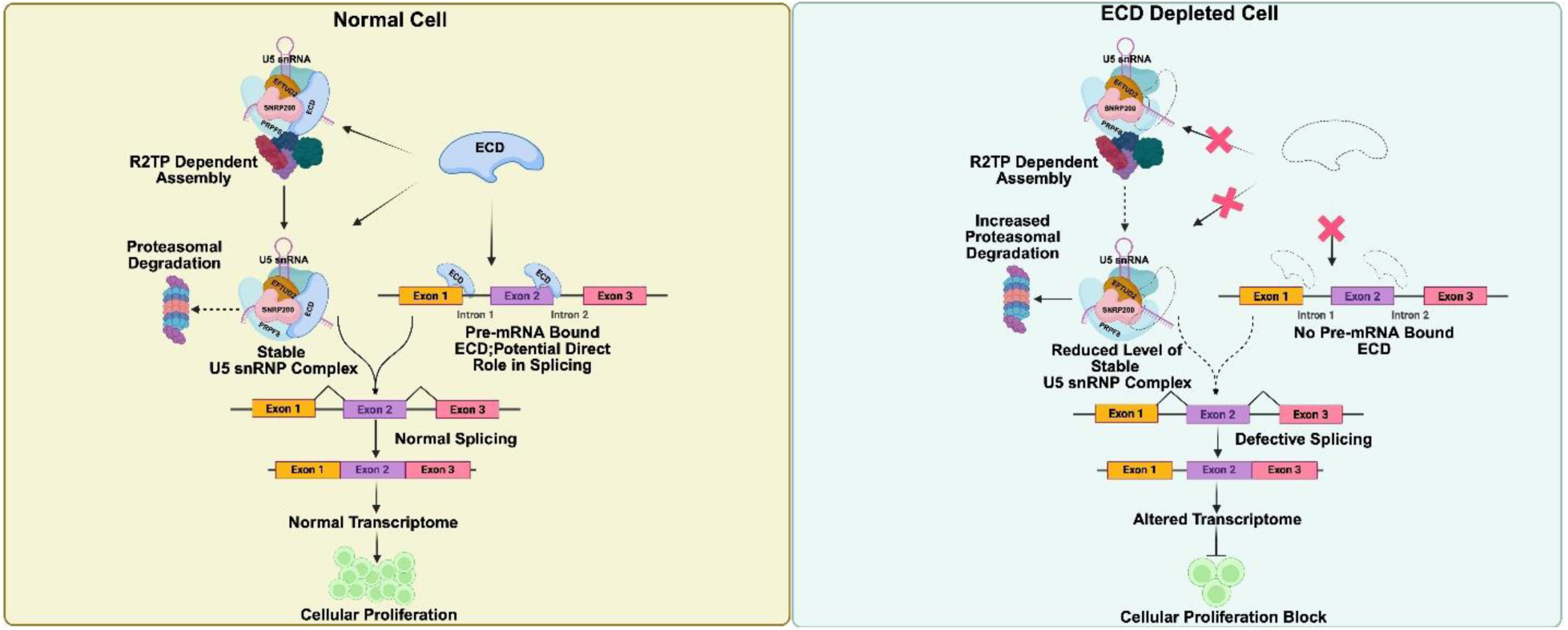

## Introduction

The interaction of cellular proteins with RNAs regulates key cellular processes, including transcription, RNA processing and translation, that are essential for cells to survive and replicate (1). RNA binding proteins (RBPs) are generally ubiquitously expressed and are among the most abundant proteins in a cell (2). Most RBPs interact with RNAs through RNA-binding domains comprised of specific amino acid sequences such as the RNA recognition motif (RRM), zinc finger (Znf) domains, heterogeneous nuclear RNP K-homology domain (KH), and others (3). However, an expanding group of noncanonical RBPs lacking the traditional RNA-binding motifs has also emerged, including transcription factors (TFs), epigenetic regulators and cell cycle regulatory proteins, significantly broadening the functional diversity of the RBPs (4). Numerous examples of such noncanonical RBPs have been characterized in the literature (5–7), where their additional RNA-binding capabilities contribute to more complex and diverse cellular functions, highlighting their multifaceted regulatory potential. In the present study, we show that ECD protein, despite lacking any traditional RNA-binding domain, directly binds to RNA and that its binding to RNA is essential for its role in RNA splicing to regulate cell proliferation.

Previously, we identified the ECD protein as a human papillomavirus HPV16 E6–binding protein (8). Subsequently, ECD has emerged as a key player in physiological processes (9–11) and various aspects of cancer development and progression (12–15). ECD overexpression in several types of cancers, including cervical and head & neck cancer (13), pancreatic cancer (16), gastric cancer (15,17), and breast cancer (18,19) is associated with poor prognosis and shorter patient survival. The co-oncogenic function of ECD was shown with mutant H-Ras for full transformation of mammary cells (12) and with HPV E7 to immortalize normal keratinocytes (13). Notably, mouse mammary epithelium-targeted transgenic overexpression of human ECD led to the development of mammary hyperplasia and tumors, demonstrating a role of overexpressed ECD in tumorigenesis (14).

Physiologically, several studies using mutational, deletional, and knockdown approaches have demonstrated the vital roles of ECD in maintaining cell survival and cell cycle progression. Mutations or deletion of *Drosophila Ecd* leads to developmental and cell survival defects (20,21). Germline deletion of the mouse *Ecd* gene led to early embryonic lethality and conditional KO of *Ecd* in *Ecd^flox/flox^* mouse embryonic fibroblasts, led to G1 cell cycle block that was rescued by ectopic human ECD expression (9). ECD’s role in cell survival was demonstrated by mitigation of ER stress by upregulating the levels of ER chaperone GRP78 (11).

More recently, ECD was found to associate with RNA biogenesis machinery proteins, such as DDX39A (19) to regulate nuclear mRNA export and with PRPF8 (19,21) to regulate mRNA splicing. Notably, *Drosophila* Ecd was found to get incorporated into and promote the assembly of the U5 small nuclear ribonucleoprotein (snRNP) complex (22). While a crystal structure of ECD has not been solved, our circular dichroism and small-angle X-ray scattering analyses suggested a folded structure of the N-terminal residues (aa 1-432) suitable for a potential scaffolding function to organize protein-protein or protein–RNA interactions (23). Given the critical role of ECD in RNA biogenesis, we examined if ECD is an RBP.

In this study, we discover and characterize ECD as a novel RBP and identify the mechanism of its role in RNA splicing through direct binding to U5 snRNA and regulating the expression and stability of U5 snRNP components. Importantly, using an RNA binding defective (RBD) mutant form of ECD, we demonstrate a requirement of RNA binding function of ECD in mRNA splicing, as well as in cell proliferation.

## Materials and Methods

### Cell lines and culture conditions

Immortalized mammary epithelial cell line, 76NTERT was cultured in DFCI-1 medium as described previously (24). *Ecd^fl/fl^*mouse embryonic fibroblasts (MEFs) were established and described previously (9), and maintained in Dulbecco’s modified Eagle’s medium (DMEM-Gibco#11965-09) supplemented with 10% fetal bovine serum (FBS-Gibco#10437-028), 10 mM HEPES (Hyclone#SH30237.01), 1 mM each of sodium pyruvate (Corning#25-000-CI), nonessential amino acids (Hyclone#SH30238.01), L-glutamine (Gibco#25030-081) and 50 μg/ml of gentamicin (Gibco#15750-078). The Lenti-X 293T cell line used for lentiviral packaging was obtained from Takara Bio (#632180) and grown in the complete DMEM medium described above. Cycloheximide (Sigma#01810) treatments were performed for various time points, as indicated in the Results Section. All cell lines were maintained in a humidified atmosphere with 5% CO2 at 37°C. Routine mycoplasma testing was performed for all the cell lines using a mycoplasma detection kit (Sigma#MP0035).

### Plasmid constructs and site-directed mutagenesis

The human ECD ORF sequence was cloned into BamHI and XhoI sites of pGEX-6p-1vector for the expression of recombinant glutathione S-transferase (GST) fusion protein of full length ECD. The GST-fused truncated mutants of ECD (1-155, 150-438, and 439-644) have been described previously (9). pGEX-6p-1 constructs carrying internal deletions within ECD (Δ135-148, Δ263–276, Δ636–644, and Δ615–625) were custom-synthesized through Gene Universal, Newark, USA. For the expression of FLAG tagged ECD in mammalian cells, a lentiviral mammalian expression vector (pLV-Puro-EF1A) carrying FLAG tag at the N-terminus of wild type (WT) human ECD was purchased from VectorBuilder Inc. Chicago, USA. A deletion mutant (Δ135-148) of ECD in the lentiviral expression vector was created using a PCR-based site-directed mutagenesis kit (NEB#E0554S), according to the manufacturer’s instructions. The primer sequences used for mutagenesis are listed in supplementary Table S1. All constructs were verified by DNA sequencing.

### Structure visualization of ECD

PDB files of predicted 3D structures of human ECD (AF-O95905-F1-v4) and *Drosophila* Ecd (AF-Q9W032-F1-v4) proteins were obtained from the AlphaFold protein structure database (25). Visualization and superimposition of structures were performed using the PyMOL tool.

### Generation of cell lines stably overexpressing WT or Δ135-148 mutant ECD

To generate cell lines constitutively expressing WT or Δ135-148 mutant ECD, lentiviral supernatants were prepared using Lenti-X 293T cells. Briefly, 293T cells were co-transfected with lentiviral packaging plasmid, psPAX2 (Addgene#12260), envelope plasmid pMD2.G (Addgene#12259), and either empty vector or vector carrying ECD (WT or mutant) using X-tremeGENE HP DNA transfection reagent (Roche#06366236001). Lentivirus containing supernatants were collected 72h post transfection. Next, 76NTERT cells were incubated with the filtered lentiviral supernatants in the presence of polybrene (8 µg/ml, Sigma#H9268). 72 hours post-transduction, cells were selected in 1 μg/ml puromycin for 7 days, and ECD levels were assessed by Western blotting (WB) using an anti-ECD monoclonal antibody (9).

### Generation of CRISPR-Cas9 mediated inducible ECD knockout cell lines, Cre recombinase mediated KO in *Ecd^fl/fl^* MEFs; siRNA mediated knockdown of PRPF8

To generate inducible ECD KO cell lines, lentiviral supernatants were prepared as described above using the Edit-R inducible lentiviral hEF1α-Blast-Cas9 plasmid (Horizon Discovery#CAS11229) and were used to transduce 76NTERT cells, followed by selection of Cas9-expressing cells in 5 μg/ml blasticidin for 7 days. These cells were then transduced with Edit-R lentiviral plasmids carrying synthetic guide RNAs against non-targeting control (NTC), (#GSG11811) or those targeting ECD: sgRNA1 (#GSGH11838-246505144; targeting sequence: TCCAAAGTCTCAACCCACAA) and sgRNA2 (#GSGH11838-246505139; targeting sequence: CTTGGGTATACTTACCCTAT). Following further selection in medium containing 5 μg/ml blasticidin and 1 μg/ml puromycin for 7 days, inducible loss of ECD in cell lines was assessed by WB after 96 hours of induction with 1μg/ml doxycycline (Sigma# D9891).

To conditionally KO Ecd in *Ecd^fl/fl^* MEFs, cells were infected with adenoviruses encoding mCherry-Cre or mCherry (control) procured from University of Iowa Viral Vector Core. 96 hours post infection, cell lysates were collected, and WB was performed to confirm the KO of *ECD*. For siRNA-mediated knockdown (KD) of PRPF8, 76NTERT cells were transfected with 40 nanomoles of control (Horizon Discovery# D-001810-10-20) or PRPF8 specific siRNA pools (Santa Cruz#sc-38209) using the DharmaFECT 1 transfection reagent (Horizon Discovery#T-2001-03) and KD verified by WB.

### Western blotting

Cell lysates were prepared in RIPA buffer, supplemented with protease (Thermo Scientific#78429) and phosphatase (Thermo Scientific#78420) inhibitor cocktails and protein concentration was measured using the bicinchoninic acid (BCA) protein assay reagent (Thermo Scientific#23227). Western blotting was performed as described previously (10,19) using the following primary antibodies: anti-ECD mouse monoclonal antibody (9,18), anti-RB (Cell Signaling#9309S), anti-p62 (Cell Signaling#5114), anti-RUVBL1 (Cell Signaling#12300S; Sigma# SAB4200194), anti-DDX39A (Abcam#ab176348), anti-PRPF8 (Abcam#ab79237), anti-SNRNP200 (Bethyl#A303-454A), anti-EFTUD2 (Abcam#ab188327), anti-PIH1D1 (Proteintech#19427-1-AP) and anti-β-actin (Sigma#A5441).

### Expression and purification of WT and mutant ECD recombinant proteins

The recombinant GST fusion proteins were expressed in bacteria and purified, as described previously (9,19). Briefly, the E. coli BL21 (DE3) transformed with the pGEX-6p-1 constructs of ECD were grown from single colonies in LB medium at 37°C to an OD600 of 0.6, followed by recombinant protein induction with 0.5 mM IPTG for 3 hours at 37°C. Bacterial lysates in sarkosyl-based lysis buffer were subjected to affinity chromatography on glutathione-Sepharose 4B beads (cytiva#17075601). The GST tag was cleaved using PreScission protease (cytiva#27084301) and removed with glutathione-Sepharose 4B beads. For His-tagged ECD protein purification, the ECD coding sequence was subcloned into the pFastBac1 vector (Thermo Scientific#10584027). Baculovirus-mediated expression in insect cells and purification of ECD using Ni-NTA affinity chromatography followed by gel filtration, were carried out by Viral Vector and Protein & Crystallography Core facilities at the University of Iowa.

### *In vitro* GST fusion protein pull-down and co-immunoprecipitation

GST fusion protein pull-downs were performed, as described previously (10,19). Briefly, 1 mg protein aliquots of 76NTERT cell lysate in Triton-X100 based lysis buffer (50 mM Tris-HCl [pH 7.5], 150 mM NaCl, 0.5% Triton X-100) were incubated with 20 μg of glutathione Sepharaose-4B bead-bound WT GST-ECD, GST-ECD △135-148 mutant or GST alone for 4 hours at 4°C. Post incubation, samples were washed five times, and bound proteins were eluted by boiling the samples in SDS PAGE sample loading buffer for 10 minutes. The bound proteins were detected by WB using the indicated antibodies. Membranes were stained with Ponceau S (Sigma#P7170) to confirm the use of comparable amounts of GST fusion proteins. For co-immunoprecipitation (co-IP) experiments, cell lysates were prepared in Triton-X100 based lysis buffer supplemented with protease and phosphatase inhibitors. For each co-IP, 1mg of protein lysate was incubated overnight at 4°C with 2.5 μg of the indicated protein specific antibodies: ECD, PRPF8, EFTUD2 or IgG antibody. Next, the protein-antibody complexes were captured by incubating the samples with protein A Sepharose beads (Thermo Fisher#101141) for an additional 3 hours at 4°C. Post incubation, samples were washed 5 times in lysis buffer and WB was performed as mentioned above. For RNase treatment, protein lysate was treated with RNase A (Thermo Fisher #EN0531) at 10 μg/mg of protein for 15 minutes at room temperature (RT).

### Electrophoretic mobility shift assay (EMSA) for ECD-RNA binding

RNA EMSA was performed to assess the direct interaction of ECD with RNA using the LightShift Chemiluminescent RNA EMSA kit (Thermo Scientific#20158). Briefly, recombinantly purified WT ECD or its various mutants were incubated with 40 nM of biotinylated RNA probes (synthesized by IDT Technologies-Probes A and B/ GenScript-Biotinylated U5 snRNA) for 45 minutes at RT in a total reaction mixture of 20 μL containing 1X RNA EMSA binding buffer (10 mM HEPES, 20 mM KCl, 1 mM MgCl2, 1 mM DTT) supplemented with 5% glycerol, 100 μg/mL tRNA (Thermo Scientific#20159) and 10 mM KCl. For antibody competition assay, ECD monoclonal antibody or control IgG was preincubated with purified ECD for 15 minutes in RNA EMSA buffer before the addition of biotinylated RNA probe. Following the incubation period, five microliters of 5× loading buffer was added to the 20 μL reaction mixture, and the samples were run on 7.5% TBE native polyacrylamide gel (pre-run at 100 V for 45 minutes in cold 0.5X TBE buffer) at 100 V until the sample dye migrated 3/4 of the length of the gel. Resolved samples were transferred to a positively charged nylon membrane (cytiva#RPN303B) in 0.5X TBE at 60 V for 60 minutes. The membranes were crosslinked with UV light using the optimal-cross link function of the Spectroline UV crosslinker instrument. Signals were visualized using the detection reagents of the biotin-labeled RNA chemiluminescence kit, as instructed by the manufacturer.

### RNA Immunoprecipitation

Immunoprecipitation (IP) of ECD-associated RNAs was performed using the IP validation and RNA imaging kit (ECLIPSEBIO#143663), as per the vendor’s instruction manual. Briefly, 76NTERT cells expressing vector or N terminal FLAG tagged ECD were UV-crosslinked at 254 nm wavelength (400 mJoules/cm^2^) using a Spectroline UV crosslinker instrument. Protein lysates from crosslinked 76NTERT cells were subjected to IP using 5 µg each of a polyclonal anti-ECD (proteintech#10192-1-AP) or a monoclonal anti-FLAG (Sigma#F1804) antibody or their respective IgG-rabbit (Sigma#PP64B) and IgG-mouse (Sigma#CS200621) control antibodies. The transcripts pulled down by the indicated antibodies were end-repaired and ligated to a biotin adapter at the 3’ end. The IP samples were run on NuPAGE 4-12% Bis-Tris protein gels, transferred to nitrocellulose membrane, and visualized using the chemiluminescent nucleic acid detection method. Concurrently, WB was performed to determine the pull-down efficiency of the antibodies. For WB, IP samples and 3% of the input samples were run on SDS-PAGE gels, transferred to PVDF membranes, and probed with the anti-ECD antibody.

### eCLIP seq and data analysis

Enhanced cross-linking and immunoprecipitation sequencing (eCLIP seq) and analyses were performed under contract by Eclipse Bioinnovations Inc, San Diego, according to the published single-end seCLIP protocol (26) with the following modifications. 20 million cells were UV crosslinked at 254 nm wavelength (400 mJoules/cm^2^) and then lysed using 1 mL of eCLIP lysis mix. Lysed samples were twice sonicated for 4 min with 30 seconds ON/OFF cycles at 75% amplitude (Q800R2 Sonicator, QSonica). 5 μg of polyclonal anti-ECD (proteintech#10192-1-AP) antibody pre-coupled to anti-rabbit IgG Dynabeads was incubated overnight with samples containing 50-100 μg of RNA at 4°C. The beads were washed with eCLIP high stringency wash buffers and collected using magnetic separation. The IP samples and 2% of paired input samples were resolved on gels, transferred to the membrane, and the region of the membrane from the ECD size to 75 kDa above was used to isolate the IPed RNA species. Subsequent RNA adapter ligation, reverse transcription, DNA adapter ligation and PCR amplification were performed as reported previously (26). Sequencing primers are ligated at 3’end and the eCLIP cDNA adapter containing a random sequence of 10 nucleotides at the 5’ end, served as a unique molecular identifier (UMI) (27). UMIs were trimmed from read sequences using the umitools (v0.5.1) (28). The UMI sequences were added to the read names in the FASTQ files to be utilized for subsequent analysis. The 3’adapters were trimmed using cutadapt (v2.7) (29) and reads shorter than 18 bp were discarded. Next, reads were mapped to human repetitive elements and rRNA sequences from Dfam (30) and Genbank (31) databases. STAR (v2.6.0c) (32) was used to map all non-repeat reads to the human genome (hg38). Utilizing UMI sequences from the read names and mapping positions, PCR duplicates were removed using Umi tools (v0.5.1). Peaks were called using CLIPper (33). For each peak, IP versus input fold enrichment scores were computed as a ratio of counts of reads overlapping the peak region in the IP and the input samples. P-value was computed for each peak by the Yates’ Chi-Square test or by the Fisher Exact test if the expected or observed read number was below 5. Different sample conditions were compared in the same manner as IP versus input enrichment. Next, for each peak (one sample type), enrichment and p-values relative to the normalized read counts overlapping these peaks in another sample type were calculated. Annotation of Peaks was done using the transcript information from GENCODE (34). To define the final annotation of the overlapping features, the priority hierarchy of CDS, UTRs, and Introns was considered, and the expected values were calculated by permutations or random peaks. Motifs from eCLIP peaks were identified using HOMER (35) and the top 10 motifs were clustered and summarized using STAMP (36). ECD eCLIP peaks were compared to peaks identified for 149 other RBPs (37), liftOver was used to convert to the hg38 genome, and an enrichment score was calculated based on the ratio of genes targeted by both the annotated RBPs and ECD vs. those targeted by ECD alone. Previously annotated functional categories were then summarized using the difference in average enrichment score for RBPs inside vs outside each category.

### Fluorescence polarization assay

Fluorescence polarization (FP) assays were performed at 25°C using a SpectraMax M5e microplate reader equipped with SoftMax Pro software (excitation at 485 nm, emission at 525 nm, cutoff at 515 nm). Purified WT or Δ135-148 mutant ECD proteins at varying concentrations were incubated with 6-FAM–labeled RNA probes containing an Sp9 linker (IDT Technologies) at a final concentration of 5 nM in binding buffer (10 mM HEPES, pH 7.4; 1 mM MgCl₂; 1 mM DTT; 50 mM KCl; 100 μg/mL BSA; 0.01% Tween-20) in black opaque 384-well plates (20 μL per well) for 30 min at room temperature in the dark. Experiments were repeated three times. FP values, expressed in millipolarization (mP) units, were plotted against the ECD protein concentration, and dissociation constants (Kd) were calculated by nonlinear regression analysis using GraphPad Prism 11 software.

### RNA-seq and mRNA Splicing Analysis

Total RNA was isolated from control (sgNTC) or ECD depleted (sgECD-2, labeled as sgECD throughout the manuscript unless specified) or control siRNA vs. PRPF8 siRNAs transfected 76NTERT cells using the Trizol reagent (Invitrogen#15596018) and the quality of purified RNA was determined using a Bioanalyzer. RNA-seq libraries were prepared using the TruSeq RNA Library Prep Kit v2 (Illumina) following the manufacturer’s protocols and subjected to 100 bp paired-end sequencing on an Illumina NovaSeq 6000 system at the UNMC Next Generation Sequencing (NGS) facility. The NGS reads from the NovaSeq 6000 server were downloaded in FASTQ format and mapped to the human genome (hg38) using STAR (v2.7.3a) (32). Differentially expressed genes (DEGs) were identified using DESeq2 (v1.44.0) (38). The normalized Transcripts Per Million (TPM) values were derived using STRINGTIE (v2.1.1), (39) and differential splicing genes (DSGs) were assessed using rmats-turbo (v4.3.0) (40). Sample-to-sample dissimilarities were calculated using Euclidean distance on the transformed data. Hierarchical clustering was then performed based on this distance matrix. The resulting dissimilarity matrix was visualized as a heatmap to assess sample relationships and clustering patterns. Function enrichment/gene ontology (GO) over-representation analysis was performed using ENRICHR (41). Signal tracks were created using deeptools (v3.5.1), (42) with BPM normalization and visualization using IGV (43). Putative transcription factor (TF) target genes were identified from the ARCHS4 database (44). The TFs within the database were then overlapped with the DSGs from this study, and the targets of identified TFs were overlapped with DEGs.

### Quantitative real-time PCR (qRT-PCR)

1µg of TRIzol-purified RNA was reverse transcribed using the iScript gDNA clear cDNA synthesis kit (Bio-Rad#1725035). qRT-PCR was conducted in a QuantStudio 3 real-time PCR system (applied biosystems) using the SYBR green master mix (Applied Biosystems#4309155). The list of primers used is provided in the Supplementary Table-S1.

### CellTiter-Glo based luminescent cell proliferation assay

*Ecd^fl/fl^* MEFs expressing the vector, WT ECD, or its △135-148 mutant were transiently infected with mCherry vector (control) or mCherry-Cre-expressing adenoviruses. 500 cells were plated per well in 96-well plates and cultured with a change of medium on alternate days. At the indicated time points, viable cells were quantified using the CellTiter-Glo 2.0 cell viability assay (Promega# G9241) according to the manufacturer’s instructions.

### Statistical analysis

The results are expressed as mean ± standard error of mean (SEM) from three independent experiments, performed at least in triplicates each time. Statistical analyses were done using the GraphPad Prism 11 software. The type of statistical test used in individual experiments is indicated in figure legends. A p-value of ≤0.05 was considered statistically significant.

## Results

### ECD directly binds to RNA through its N-terminal region

In previously published analyses of ECD-associated proteome, we identified multiple proteins involved in RNA biogenesis, including the mRNA splicing-associated, U5 small nuclear ribonuclear protein 200 (SNRNP200), elongation factor Tu GTP binding domain containing 2 (EFTUD2), spliceosome scaffold protein PRPF8, and the mRNA export protein DDX39A (19). Additionally, *Drosophila* Ecd interaction with Prp8 (ortholog of mammalian PRPF8) was shown to regulate alternative mRNA splicing (21) through Prp8 stability by Ecd leading to the promotion of U5 snRNP maturation (22). Given the critical role of ECD in RNA biogenesis, we examined if ECD directly bind to RNA. The online tool RNABindRPlus (45) predicted three regions in ECD (amino acids (aa) 263-276, aa 615-625, and aa 636-644) with high scores for potential RNA binding **(Fig. 1A)**. Additionally, three other sites (aa 135–148, aa 205-219, and aa 301-309) in the N-terminal region were identified with lower prediction scores (**Fig. 1A).** We therefore examined the RNA binding potential of ECD by performing RNA EMSA using purified ECD protein and a biotinylated single-stranded RNA probe, previously reported to bind DDX39A (designated as Probe-A in this study) (46). The GST tag of the E. coli-purified recombinant ECD fusion protein was cleaved with precision protease, and purity was confirmed by Coomassie staining of SDS-PAGE gels **(Fig. S1A)**. The RNA EMSA showed an ECD dose-dependent upward shift in the migration of the RNA probe signifying the formation of an ECD-RNA complex **(Fig. 1B)**. The specificity of binding was demonstrated by efficient competition by an excess amount of the unlabeled RNA probe **(Fig. 1B)**. Further, inclusion of an ECD specific monoclonal antibody eliminated the ECD RNA binding while the IgG control had no effect **(Fig. 1C)**. To further validate these results, we generated His-tagged ECD protein using a baculovirus system. Comparable purity was observed for GST-affinity purified **(Fig. S1A)** and Ni-NTA affinity purified His tagged ECD protein **(Fig. S1B),** and the RNA EMSA results with the His tagged ECD protein (**Fig. S1C)** were comparable to those observed with the GST-affinity purified ECD protein (**Fig. 1B)**, further supporting our findings. To localize the region(s) needed for RNA binding, we conducted EMSA assays with GST-purified deletion fragments of ECD that included the N-terminal (aa 1-155), middle (aa 150-438), and C-terminal (aa 439-644) **(Fig. S1D)**. These analyses revealed that only the N-terminal region (aa 1-155) of ECD binds to RNA **(Fig. 1D)**. Interestingly, the RNABindRPlus predicted ECD N-terminal region (aa 135–148), though having a low score for RNA binding was the only predicted site within the aa 1-155 region **(Fig. 1A)**. Thus, to further validate our findings, we purified ECD deletion mutants, either lacking aa 135-148 (ECDΔ135–148) or other top three predicted amino acid regions (aa 263–276, 615–625 and 636–644) using GST-affinity chromatography (**Fig. S1E)**. Notably, only ECDΔ135-148 mutant exhibited significant decrease in RNA binding **(Fig. 1E),** substantiating the conclusion that the N-terminal region of ECD binds to RNA and that aa 135-148 contribute to this activity. Next, we conducted an independent assay of RNA binding using a fluorescence polarization (FP) assay and compared the binding affinities of WT and ECDΔ135–148 for the RNA probe A. Notably, the ECDΔ135–148 protein showed a two-fold increase in the dissociation constant (Kd) value (9.3 ± 2.5 µM) relative to that with the WT protein (4.5 ± 0.9 µM) **(Fig. 1F).** Importantly, the AlphaFold-predicted structure of human ECD shows that aa 135-148 are part of anti-parallel beta sheets **(Fig. 1G),** and the corresponding region (aa 163–173) in *Drosophila* Ecd adopts an identical secondary structure **(Fig. S2).** Consistent with a biologically key role of the RNA-binding region of ECD, the AlphaMisfold tool predicts high pathogenicity scores for amino acids around this region **(Fig. 1H).** Anti-parallel beta-sheet structures are well-documented in literature as potential RNA-binding domains (3). Together, our findings provide compelling evidence that ECD directly interacts with RNA through its N-terminal region and support the notion that RNA binding by ECD may underlie its biological roles.

**Fig. 1.**
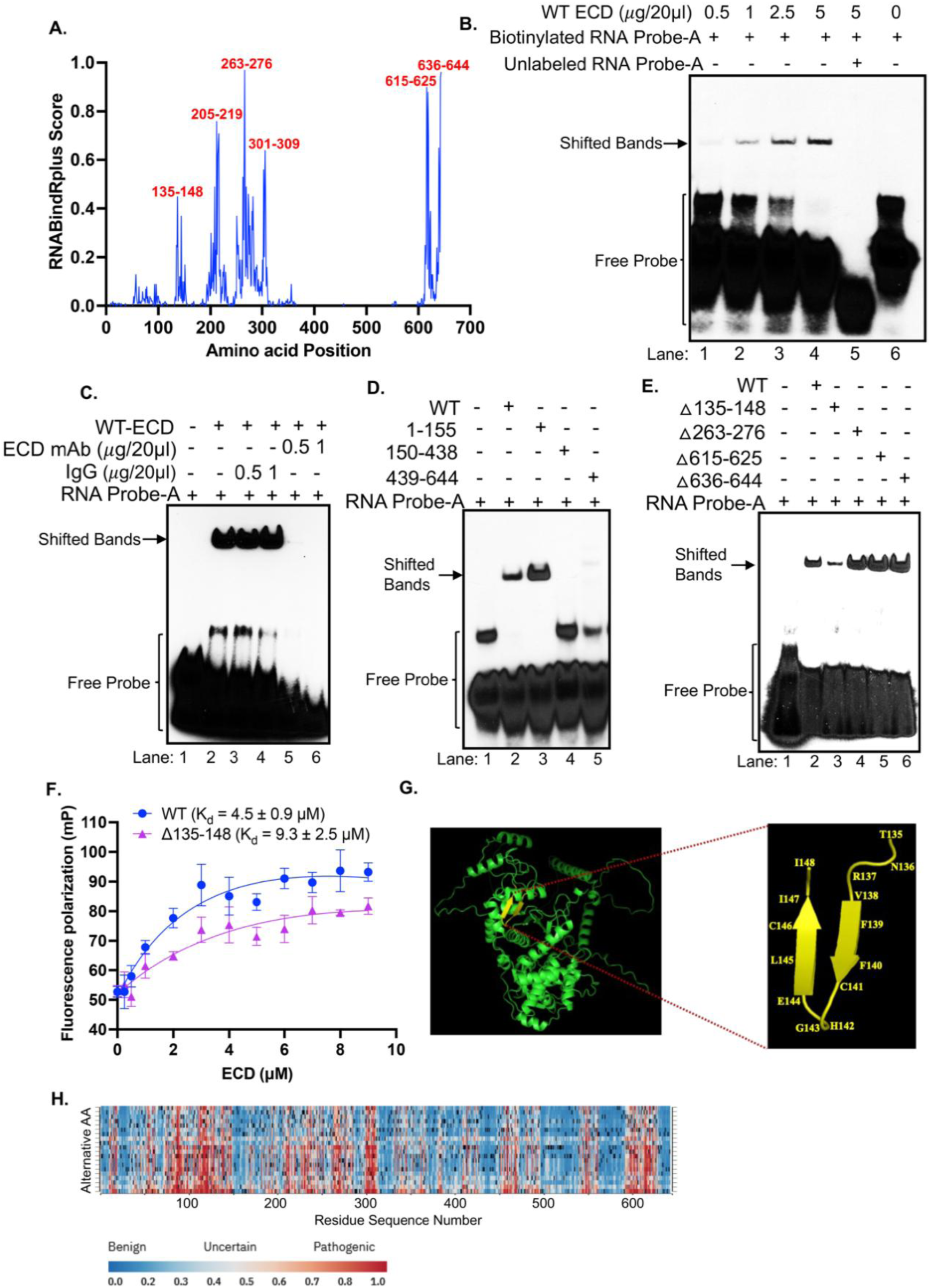
ECD directly binds to RNA via the N-terminal region. **(A)** ECD amino acids predicted to bind RNA using online tool RNABindRplus. **(B)** RNA-EMSA with different concentrations of purified ECD (lanes 1 to 4) along with biotinylated RNA probe (40 nM) previously reported to bind DDX39A (Probe-A). Protein free probe, used as a negative control (lane 6). Competition with 100-fold excess unlabeled probe was done to determine the binding specificity (lane 5). **(C)** RNA EMSA with purified ECD (2.5 µg/20 µl), with RNA Probe-A (all lanes), in the absence (lane 2) or presence of two different concentrations of mouse ECD monoclonal antibody (lanes 5 and 6) or equal amount of mouse IgG (lanes 3 and 4). Protein free probe served as negative control (lane 1). **(D)** RNA-EMSA with 2.5 µg /20 µl of wild type (WT, lane 1) or same amount of (2.5 µg/20 µl) indicated truncated mutants of ECD (lanes 2 to 5) along with Probe-A. **(E)** Purified deletion mutants used in RNA EMSA (2.5 µg/20 µl) show loss of RNA binding upon deletion of amino acids (aa) 135 to 148 (lane 3) while WT (lane 1) and other deletion mutants of ECD (lanes 4 to 6) show binding to RNA. All EMSA samples were run on 7.5% native gel, transferred to Hybond-N+ membrane, and developed with Streptavidin-HRP based chemiluminescent method. **(F)** Binding affinities of WT and Δ135-148 mutant of ECD to RNA probe-A as determined by fluorescence polarization assay. **(G)** Predicted tertiary structures of human ECD, showing identified RNA binding region, aa 135-148 (in yellow). PDB file of Alpha Fold predicted 3D structure of human ECD was visualized using PyMOL software. **(H)** Alpha Misfold prediction of mutation pathogenicity per residue of human ECD.

### ECD binds to cellular RNA

To assess the ECD’s RNA binding ability in cells, we performed UV crosslinking and immunoprecipitation (CLIP) using 76NTERT cells stably expressing either vector or FLAG-tagged ECD. RNA species migrating over a large range on gels were detected in anti-FLAG IPs of FLAG-ECD-expressing but not in vector cells, and such species were detected in with or without FLAG-tagged expressing cell lines with anti-ECD antibody, with more robust RNA-IP in FLAG-ECD expressing cells (**Fig. 2A**). As expected, control mouse or rabbit IgG antibody IPs did not show any co-IPed RNAs **(Fig. 2A).** Concurrent anti-FLAG **(Fig. 2B)** or anti-ECD **(Fig. 2C)** immunoblotting demonstrated the efficiency of IPs and lack of pull-down with respective control IgGs further assures the specificity. These results clearly support the notion that ECD binds to RNA within the cell.

**Fig. 2.**
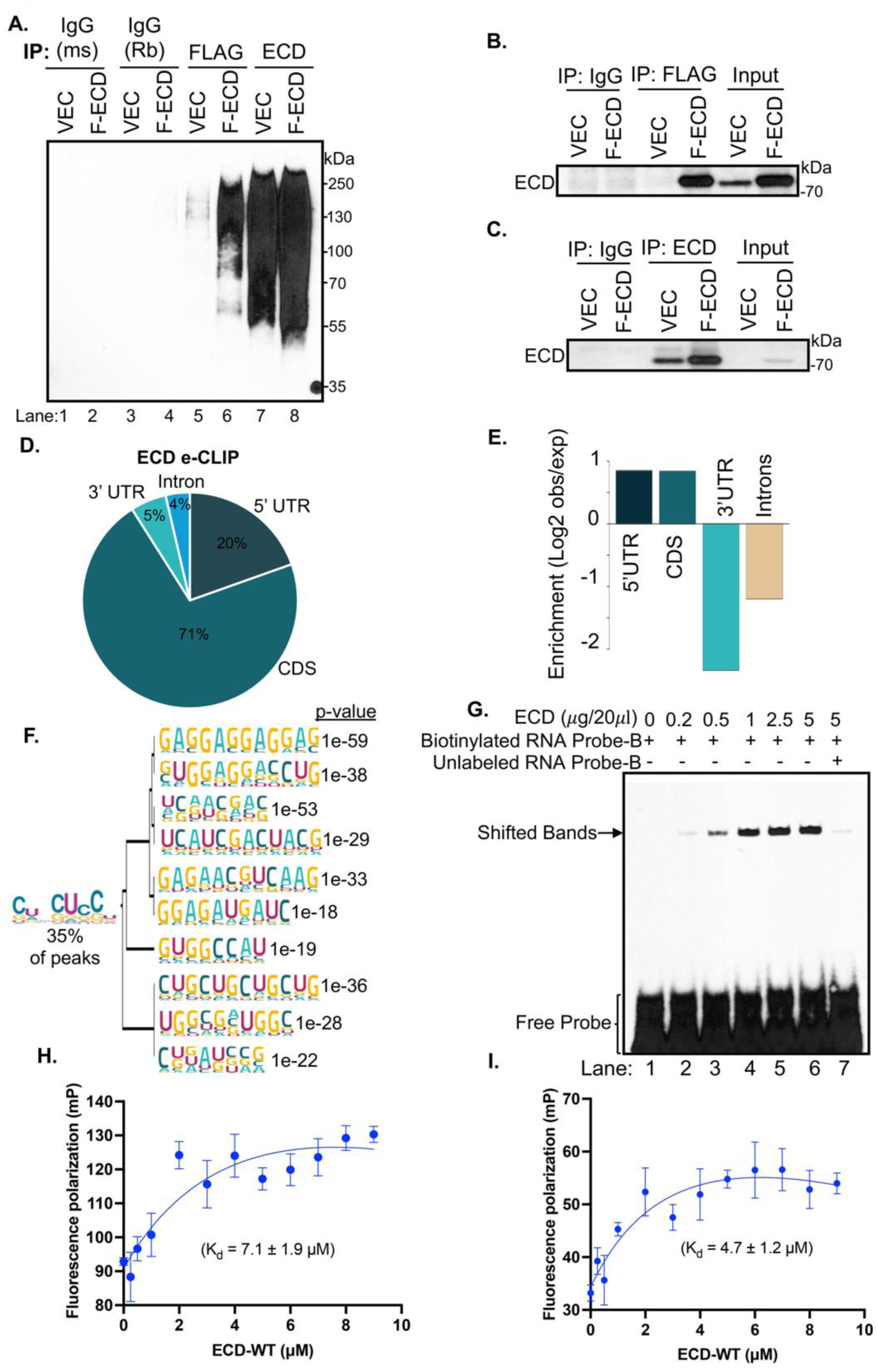
RNA binding landscape of ECD by eCLIP seq. (**A)** Blot depicting immunoprecipitated (IP) RNA. Protein lysates from UV crosslinked 76NTERT cells expressing vector or FLAG-ECD were subjected to IP using 5 µg each of anti-ECD or FLAG or respective IgG (rabbit-Rb) and IgG (mouse-ms) antibodies. The transcripts pull down by indicated antibodies were end repaired and ligated with a biotin adapter at 3’ end as per the protocol by ECLIPSEBIO antibody IP validation kit. IP samples were run on NuPAGE 4-12% Bis-Tris protein gels, transferred to nitrocellulose membrane, and visualized using the chemiluminescent nucleic acid detection method. **(B&C)** WB depicting IPed ECD protein using anti-FLAG **(B)** and anti-ECD **(C)** antibodies. For WB, IP and 3% of input samples were run on SDS-PAGE gels and transferred to PVDF membrane and probed with ECD antibody. **(D)** The distribution profile of the ECD eCLIP peaks in the human genome from biological duplicates. CDS-coding sequence, UTR-untranslated region**. (E)** Enrichment of ECD eCLIP-seq peaks compared to random distribution (expected) to account for the different frequency and size of individual elements. **(F)** The top-ranked motifs discovered de novo from ECD eCLIP-seq peaks by HOMER analysis along with a consensus motif (present in 35% of ECD eCLIP peaks) identified through clustering of top 10 HOMER motifs using STAMP. **(G)** RNA-EMSA with indicated concentrations of purified ECD (lanes 2 to 4) along with 40 nM of biotinylated RNA probe designed with eCLIP identified the third-ranked purine-pyrimidine-mix motif UCAACGAC (Probe-B). Protein free probe, used as a negative control (lane 1). Competition with 100-fold excess unlabeled probe was performed to determine the binding specificity (lane 7). Samples were processed as mentioned in Fig. 1. **(H&I)** Fluorescence polarization binding curves depicting the binding affinity of WT ECD to RNA probes designed from eCLIP-identified motifs. **(H)** RNA probe comprising the top-ranked purine-only GAG repeat motif, and **(I)** RNA probe containing the third-ranked purine-pyrimidine-mix UCAACGAC motif.

Next, to directly assess cellular RNAs associated with ECD under physiological conditions, we performed enhanced UV crosslinking and IP followed by sequencing (eCLIP-seq) analysis (47) using an anti-ECD polyclonal antibody to IP ECD-associated RNAs in 76NTERT cells. The eCLIP-seq analysis revealed a total of 3,363 transcriptome-wide ECD binding sites that were shared between replicates based on irreproducible discovery rate (IDR) analysis using the CLIPper pipeline (33). A vast majority of ECD binding sites were primarily localized in coding exons (CDSs-71%), and 5′ untranslated regions (5’UTRs-20%) with minimal overlap with 3′UTRs (5%), and introns (4%) **(Fig. 2D)**. Monte Carlo permutation testing to control for relative feature sizes showed enrichment of ECD in both CDSs and 5’UTRs **(Fig. 2E).** De novo analysis of peak-associated motifs identified several significant sequences, summarized as GA(UC) rich sequences **(Fig. 2F)**. Notably, only 35% of ECD peaks overlapped these motifs, indicating that while ECD may prefer GA(UC) rich sequences, binding to RNA is not associated with one specific sequence **(Fig. 2F)**. An EMSA using a biotinylated RNA probe comprising one of the eCLIP-identified RNA motifs—the third motif from the top, a mixed purine–pyrimidine sequence (UCAACGAC), confirmed that the recombinantly purified ECD binds eCLIP identified RNA sequences. Binding was efficiently competed by excess unlabeled probe **(Fig. 2G).** Next, we assessed the ability of ECD to bind to specific RNA sequences identified by eCLIP. We performed a FP assay using fluorescent RNA probes comprising the top motif, a purine-exclusive GAG repeat **(Fig. 2F),** and the third purine–pyrimidine mixed motif (UCAACGAC). The results showed that ECD binds the purine-exclusive motif with slightly lower affinity with a dissociation constant of 7.1 ± 1.9 µM **(Fig. 2H)** than the mixed motif (dissociation constant of 4.7 ± 1.2 µM) **(Fig. 2I).** These results corroborate the eCLIP-seq findings, further solidifying the conclusion that ECD functions as an RNA-binding protein.

GO biological process pathway analysis of ECD binding RNAs that possess the identified ECD-binding consensus motif showed enrichment for key cellular pathways including the transcription, apoptosis, response to endoplasmic reticulum stress and cell proliferation **(Fig. S3A).** These pathways play a crucial role in physiological functions and are commonly altered in oncogenesis and tumor progression, consistent with previously identified roles of ECD in physiological context (9,11) and tumor initiation and progression (14,16).

### ECD deficiency leads to transcriptome-wide alteration in gene expression and causes aberrant retention of spliced-out RNAs

Given the broad repertoire of cellular mRNAs that we identified to be in complex with ECD, we performed RNA-seq analysis of control vs. ECD depleted 76NTERT cells using doxycycline-regulated Cas9 for short-term ECD loss since stable KO is not compatible with cell viability (9). Before RNA-seq analysis, loss of ECD was checked concurrently in cell lysates from doxycycline treated control (sgNTC) and sgECD groups. Data revealed a marked reduction in ECD protein levels upon doxycycline-induced depletion **(Fig. 3A).** RNA-seq analysis revealed that the replicates of both the control and ECD-depleted conditions were similar within their respective groups **(Fig. S3B)** and ECD depleted group showed distinct transcriptional patterns from control cells **(Fig. 3B).** Differential expression testing using log2 FC ≥ 1 and adjusted P-value < 0.05 as a cutoff, showed differential expression of 3,452 genes, with 1,317 downregulated and 2,135 upregulated genes in ECD depleted cells **(Fig. 3C).** Biotype analysis revealed that upon ECD depletion, DEGs corresponded to protein coding (34.4%), lncRNA (29.2%) and Pseudogene (26.4%) categories **(Fig. S3C).** MSigDB gene set analysis of DEGs showed that those downregulated upon ECD depletion are enriched in cholesterol homeostasis, E2F target, KRAS signaling, and G2-M checkpoint pathways, whereas the upregulated genes are enriched in inflammatory response, TNF-alpha and hypoxia signaling pathways **(Fig. 3D).** GO biological process analysis also indicated an enrichment of downregulated genes in sterol and cholesterol biosynthetic processes along with protein-DNA and nucleosome assembly, while upregulated genes corresponded to regulation of type II interferon production **(Fig. 3E).** Heatmaps illustrate the downregulation of cholesterol homeostasis and protein-DNA complex assembly related genes **(Fig. 3F)** and upregulation of inflammatory response and negative regulation of type II interferon pathway genes **(Fig. 3G)** upon ECD depletion. These results support a key role of ECD in ensuring the expression of multiple gene networks involved in diverse physiological functions.

**Fig. 3.**
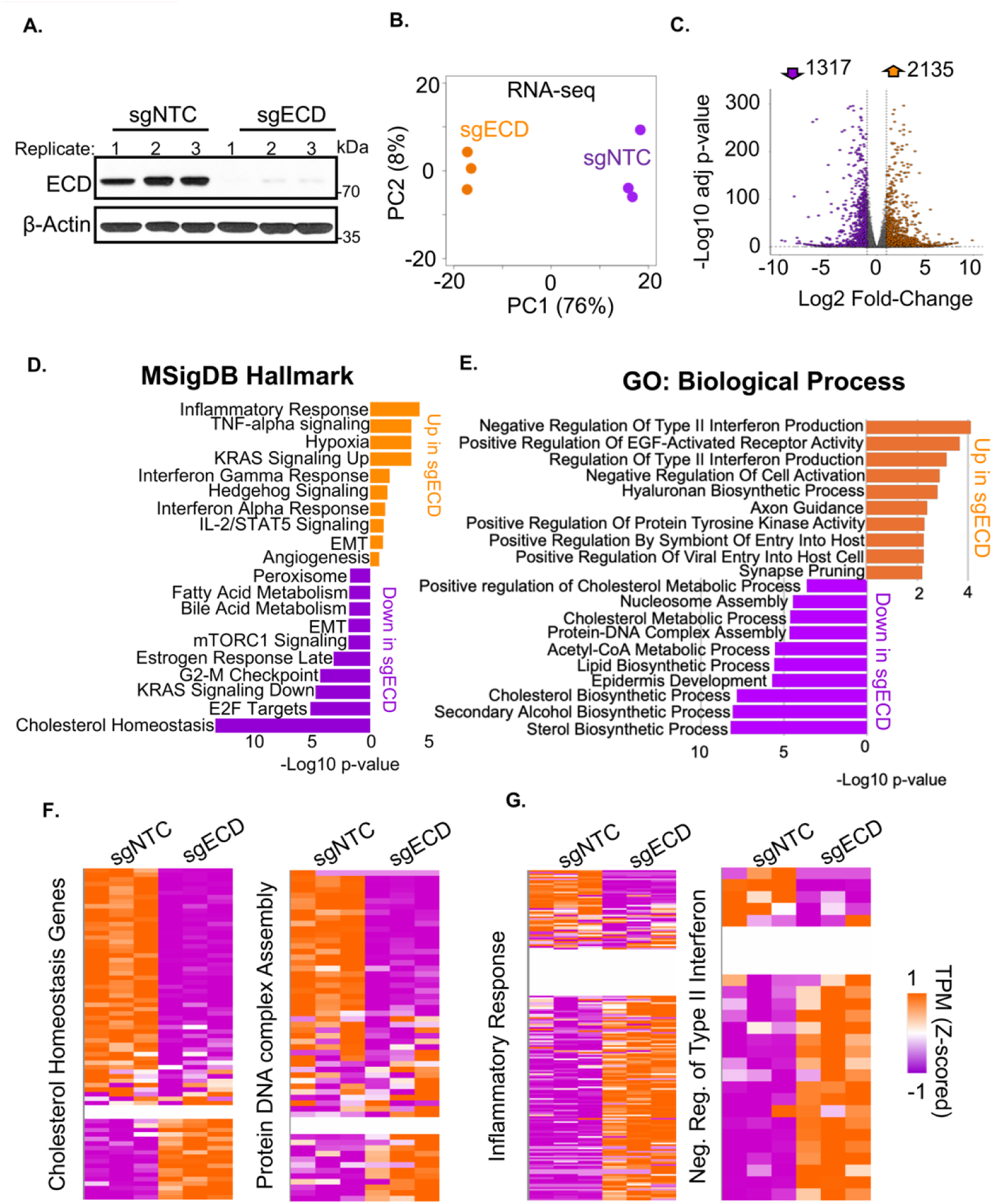
ECD depletion alters transcriptomic profile. **(A)** WB depicting expression of ECD protein in the control (sgNTC) and ECD depleted (sgECD) 76NTERT cells used for RNA seq analysis, following doxycycline-induced Cas9 activation. β-Actin used as loading control. **(B)** PCA of RNA-seq data done in biological triplicates after ECD depletion (orange) compared to non-targeting controls (sgNTC, purple). **(C)** Volcano plot of differential expression after ECD depletion identified thousands (1,317 decreased and 2,135 increased) of significant differentially expressed genes (Benjamini-Hochberd corrected p-value < 0.05 and >2-fold change). **(D-E)** The top 10 increased and decreased MsigDB Hallmark pathways and Gene Ontology Molecular Function gene sets associated with differential expression post ECD depletion. **(F-G)** Heatmaps of the z-scored expression (Transcripts Per Million) for the specified gene sets.

Given our (13) and others’ (21,22) published studies linking ECD to regulation of mRNA splicing, we evaluated the impact of ECD depletion on the transcriptome-wide mRNA splicing. Based on an FDR of < .05 cutoff, we observed differential splicing pattern in ECD depleted cells **(Fig. 4A).** The rMATs tool identified a total of 5,070 DSEs in ECD depleted cells as compared to control cells and these splicing events map to 2,595 unique genes, suggesting many genes had multiple differential splice events. Indeed, 1,844 genes showed one, 527 genes showed two, 141 genes showed three, and 83 genes showed four or more DSEs **(Fig. 4B).** Notably, exon skipping and intron retention were the most altered splicing events upon ECD depletion, while the impact on other splicing events like alternative 5’ splice site and 3’splice site usage was less **(Fig. 4C).** ECD depleted cells showed downregulation at 1,373 exon skipping events out of the 5,070 total DSEs. In contrast, aberrant splicing involving intron retention was amplified, with 2,220 intron retention events out of 5,070 total identified DSEs **(Fig. 4C).** Interestingly, MSigDB pathway analysis showed an enrichment for mitotic spindle and DNA repair pathways among genes exhibiting altered splicing upon ECD depletion **(Fig. 4D).** GO molecular function analysis revealed enrichment for RNA binding and cadherin binding functions for genes with altered splicing upon ECD depletion **(Fig. 4E).** Furthermore, GO biological process revealed enrichment for transcription and chromatin modeling pathways among the DSGs observed upon ECD depletion **(Fig. S4A).** RNA-seq signal tracks of representative genes belonging to top altered categories, *ARAP3* from mitotic spindle **(Fig. 4F)**, *MOV10* from RNA binding **(Fig. 4G),** and *CCNT2* from positive regulation of DNA templated transcription categories **(Fig. S4B)** illustrating aberrant RNA retention after ECD depletion. Overall, these results support the importance of ECD in regulating transcriptome-wide mRNA splicing events and transcript levels in mammalian cells to help maintain diverse and functionally critical molecular pathways.

**Fig. 4.**
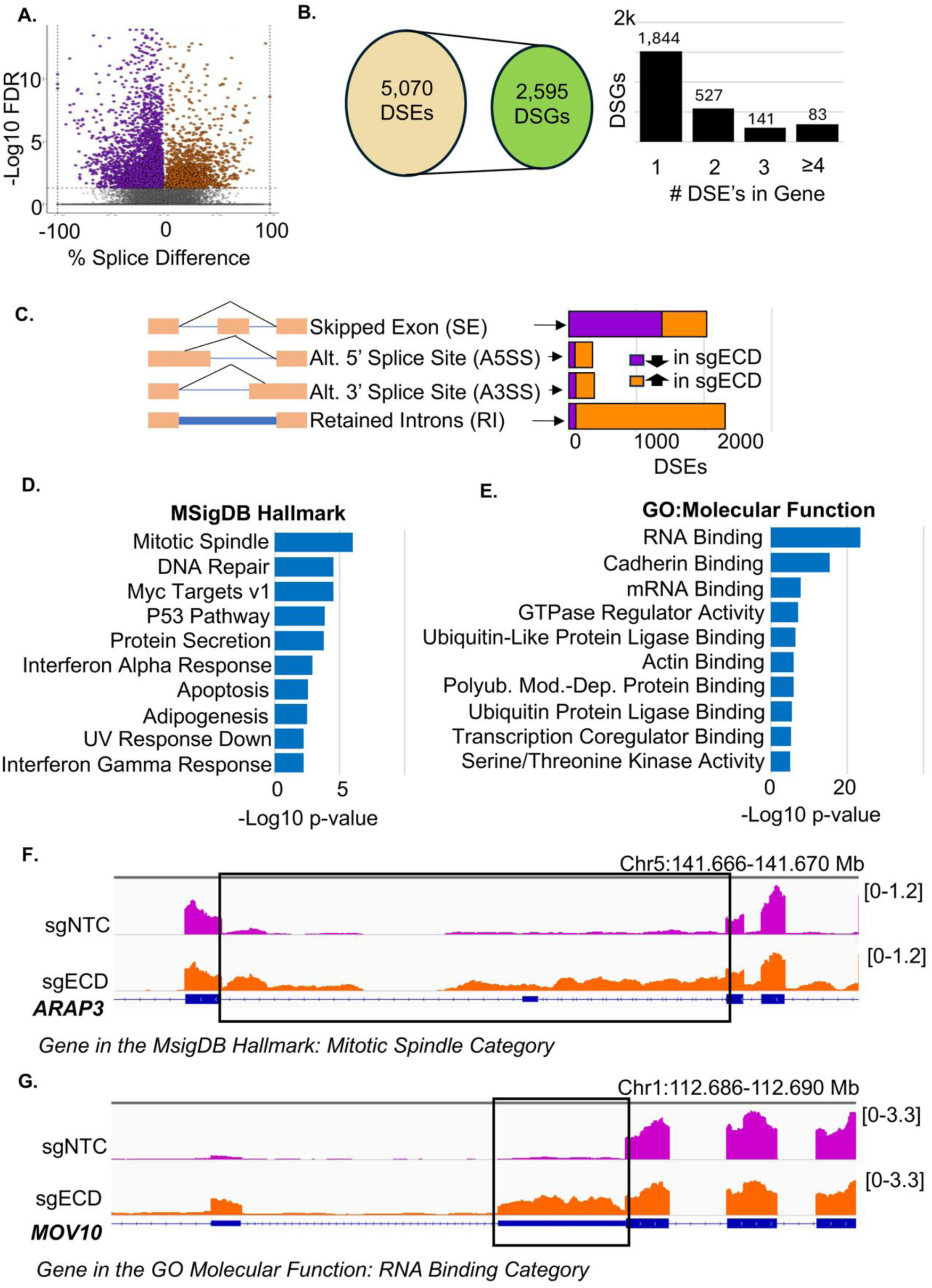
ECD expression impacts RNA splicing. **(A)** Results of differential splicing analysis, which identified thousands of significant (FDR < 0.05) differential splice events (DSEs). Schematic and Bar graph showing that: **(B)** many genes have multiple differential splice events (e.g. multiple retained introns), such that 5,070 differential splice events are mapped to 2,595 genes; and **(C)** categorization of differential splice events identified primarily aberrant increased retention of features (an increase of retained introns and a decrease of skipped exons). **(D-E)** The top MsigDB Hallmark pathways and Gene Ontology Molecular Function gene sets associated with differentially spliced genes upon ECD loss. **(F-G)** RNA-seq signal tracks showing aberrant retention in transcripts of genes (in the top categories shown in D and E) post ECD depletion.

### ECD binding to RNAs is associated with regulation of alternative splicing

To delineate if the RNA-binding activity of ECD is linked to its role in mRNA splicing and/or gene expression, we first analyzed the overlap between DEGs or DSGs upon ECD depletion. We found minimal overlap between 3,452 DEGs and 2,595 DSGs, with only 179 genes shared between the two groups **(Fig. 5A).** To probe this relationship more directly, we assessed the overlap of ECD eCLIP peaks with DEGs and DSGs vs. randomly selected negative control genes. While only 5.56% of DEGs overlapped with ECD eCLIP peaks, a significantly higher proportion, 22.19%, of DSGs showed such overlap **(Fig. 5B)**. Additionally, mapping the average profile of ECD eCLIP enrichment signals (Log2 IP/input) at differentially spliced coordinates revealed enhanced enrichment signals at mis-spliced junctions **(Fig. 5C)**. Visualization of representative gene tracks, *AGRN* **(Fig. 5D)** and *MOV10* **(Fig. 5E)**, demonstrates the ECD occupancy on transcripts that undergo mis-splicing following ECD depletion. We observed the ECD binding near splice sites as well as a diffuse distribution across the transcript. Collectively, these findings support the idea that ECD interacts with pre-mRNAs and regulates alternative splicing events.

**Fig. 5.**
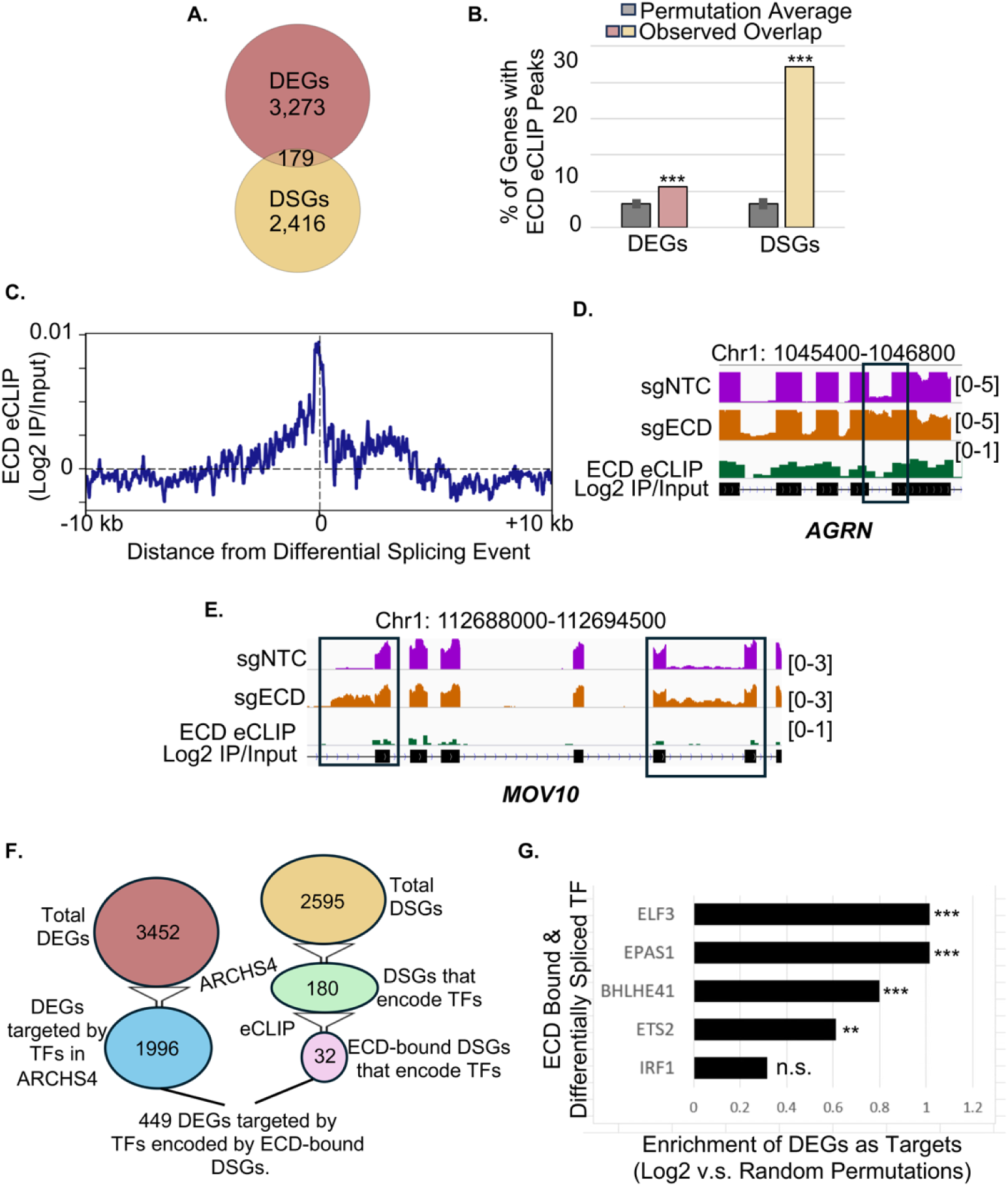
ECD binds RNA to modulate splicing. (**A)** Venn diagram depicting, differentially expressed genes (DEGs) and differentially spliced genes (DSGs) have minimal overlap **(B)** Bar graphs showing percentage of DEGs and DSGs that overlap an ECD e-CLIP peak. *** indicate p-value < 0.001 (Monte Carlo permutation testing) compared to the average overlap of peaks with randomly selected genes (grey) with error bars representing the standard deviation from a 1,000 permutation tests. **(C)** Average profile of ECD e-CLIP enrichment signal (Log2 IP/input) at differentially spliced coordinates. **(D & E)** Examples of RNA-seq signal at introns that are retained after ECD depletion. Shown below is the ECD e-CLIP signal in the control cells. **(F)** Schematic of the filtering pipeline to identify gene targets as ECD bound transcripts that are differential spliced and encode transcription factors (TFs) that putatively regulate differentially expressed genes in the ARCHS4 database. **(G)** Top 5 TFs based on the number of DEGs that they regulate relative to randomly selected genes. *** p<0.001, ** p<0.01, ns=non-significant, computed through Monte Carlo permutation testing.

The weaker linkage of ECD’s RNA binding activity with DEGs **(Fig. 5B)** suggested the likelihood that regulation of gene expression by ECD may not be through direct association with RNAs. We therefore speculated that ECD regulates gene expression indirectly by altering the splicing of genes that may encode transcriptional factors (TFs), affecting their target genes. Consistent with this speculation, out of 2,595 identified DSGs, 180 genes were found to encode TFs, and 32 of these were bound to ECD in our eCLIP-seq analyses. Concurrently, we analyzed the ARCHS4 database and identified 1,996 genes exhibiting differential expression upon ECD loss as direct targets of TFs in this database. Among these, 449 genes were regulated by TFs whose RNAs were directly bound to ECD and aberrantly spliced. **(Fig. 5F).** Ranking of TFs whose RNAs were bound to ECD and differentially spliced upon ECD depletion based on the number of their targets exhibiting differential expression (compared to randomly selected genes) identified four TFs, ELF3, EPAS1, BHLHE41, and IRF1 that showed significant enrichment of DEGs among their targets **(Fig. 5G).** These results suggest that ECD binds and splices RNAs of TFs to regulate the expression of diverse genes.

Next, we examined how the RNA repertoire bound by ECD compares with that of other well-established RNA-binding proteins to develop a comprehensive understanding of sequence specificity of RNAs bound to ECD and to do so we compared ECD eCLIP signal to that of 149 various RBPs from a reference eCLIP dataset (37). Interestingly, based on overlap enrichment ratio, the top 10 RBPs with RNA-binding profiles similar to ECD included PRPF8 and EFTUD2 **(Fig. 6A),** two U5 snRNP complex proteins which we found previously to interact with ECD (19). We then examined previously annotated functional categories that were assigned to RBPs and, notably, “spliceosome” and “splicing regulation” emerged as the top-hits for RBPs that have similar RNA binding profiles to that of ECD **(Fig. 6B)**. Moreover, the average eCLIP signal plots for each of the top 10 spliceosome-associated RBPs around ECD dependent differential splicing events demonstrated that these proteins bind to RNA near sites that become mis-spliced in the ECD depleted cells **(Fig. 6C)**. These findings reinforce the potential involvement of ECD, bound to mRNAs, in RNA processing and splicing-related mechanisms. Additionally, as an example of the overlap, we compared the eCLIP signal tracks for each of the top 10 RBPs within the spliceosome category at *TMEM214*, a representative gene that exhibits both ECD eCLIP signal and increased intron retention in ECD depleted cells. As observed earlier with gene tracks of *AGRN* and *MOV10*, our analysis revealed that ECD displays a diffuse binding signal across the gene rather than a sharply defined single binding site, like other RBPs annotated in the spliceosome category **(Fig. 6D)**. These observations argue against a strictly sequence-driven binding model for ECD. Overall, these findings suggest a potential direct role for ECD in modulating RNA splicing, possibly in cooperation with other splicing factors, for example through its continued requirement for maintaining the stability of the U5 snRNP during splicing. In-depth studies of mRNA splicing and structural studies of spliceosome in wildtype and ECD-depleted cell models are needed to test these suggestions.

**Fig. 6.**
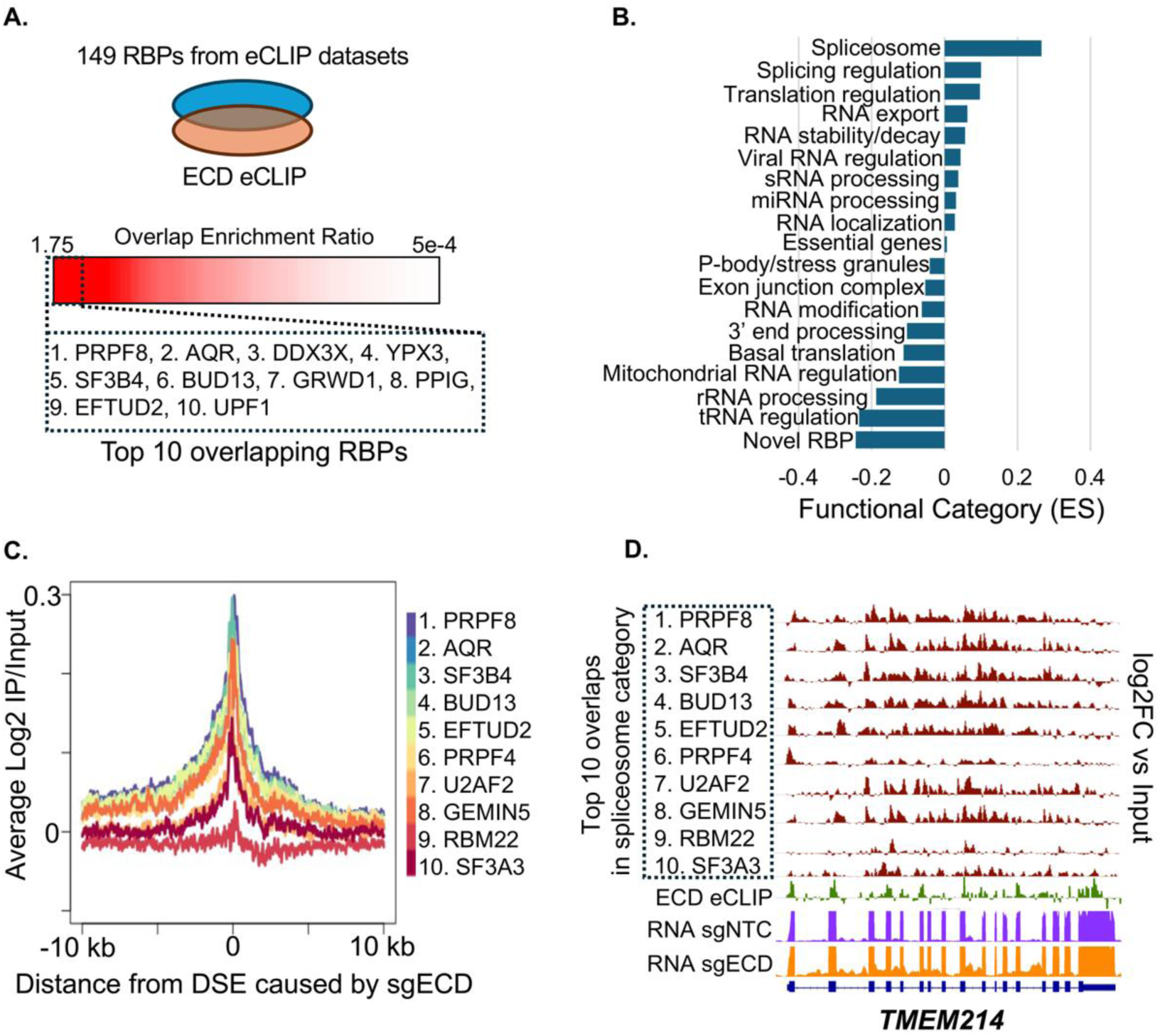
ECD bound transcriptomic landscape overlaps with RNA binding proteins involved in splicing regulation. **(A)** 149 RBPs with eCLIP data were tested for the overlap of targets with that of ECD eCLIP. Overlap enrichment ratio is calculated as the fraction of ECD peaks that overlap the RBP peaks vs those that don’t. The top 10 RBPs (ranked by overlap enrichment) are listed. **(B)** The relative enrichment of functional categories associated with RBPs that overlap ECD eCLIP targets. (ES-enrichment score). **(C)** Average signal profiles of top 10 spliceosome related RBPs centered around differential splice events caused by sgECD. **(D)** eCLIP signals at an example gene, *TMEM214* for each of the top 10 RBPs in the spliceosome category and ECD.

### ECD directly interacts with U5 snRNA and maintains the expression of key components of the U5 snRNP complex

Our results that RNAs of only ∼22% of DSGs in ECD depleted cells are directly bound to ECD, suggested additional mechanisms by which ECD regulates RNA splicing. Since we have shown that ECD associates with proteins that are components of the U5 snRNP complex (19) and others demonstrated that *Drosophila* Ecd regulates the stability of the U5 component Prp8 and also co-IPs with U5 snRNA (22), we assessed if the RNA binding activity of ECD may be linked to the regulation of U5 snRNP. First, we examined the impact of ECD depletion in cells on the levels of key U5-specific proteins. As reported, ECD depletion not only led to the downregulation of PRPF8 levels (22) but also led to a marked reduction in the levels of other U5-specific proteins, SNRNP200 and EFTUD2 **(Fig. 7A)**. To explore the mechanism underlying this downregulation, we performed the cycloheximide (CHX) chase analyses in control and ECD-depleted cells. CHX treatment revealed that ECD depletion reduces the stability of the U5 snRNP components PRPF8, EFTUD2, and SNRNP200 **(Fig. 7B-E)**. Since ECD is known to interact with components of the R2TP complex (10,48), a co-chaperone involved in early phases of U5 snRNP assembly and maturation, we investigated whether the instability of U5-specific proteins upon ECD depletion altered their interaction with the R2TP complex. To test this, we performed co-immunoprecipitation analyses to assess the association between the R2TP components and U5-specific proteins in control and ECD depleted cells. Notably, despite reduced levels of U5 proteins in ECD-depleted cells, we found more PRPF8 and EFTUD2 to be associated with the R2TP complex components, RUVBL1 and PIH1D1 (R2TP components known to interact with ECD) (10), suggesting that the absence of ECD protein contributes to a more persistent association of the U5 proteins with the R2TP complex **(Fig. 7F-J)**. In a complementary approach, to account for the reduced immunoprecipitation of U5 proteins in ECD depleted conditions, we used 1.4-fold more protein lysate for PRPF8 and EFTUD2 immunoprecipitations in ECD-depleted cells as compared to the control. This approach further confirmed an increased interaction of the R2TP components RUVBL1 and PIH1D1 with U5 proteins PRPF8 and EFTUD2 **(Fig. S5A-E).** Taking together, these results suggest ECD depletion results in a more persistent association of U5-specific proteins with the R2TP complex.

**Fig. 7.**
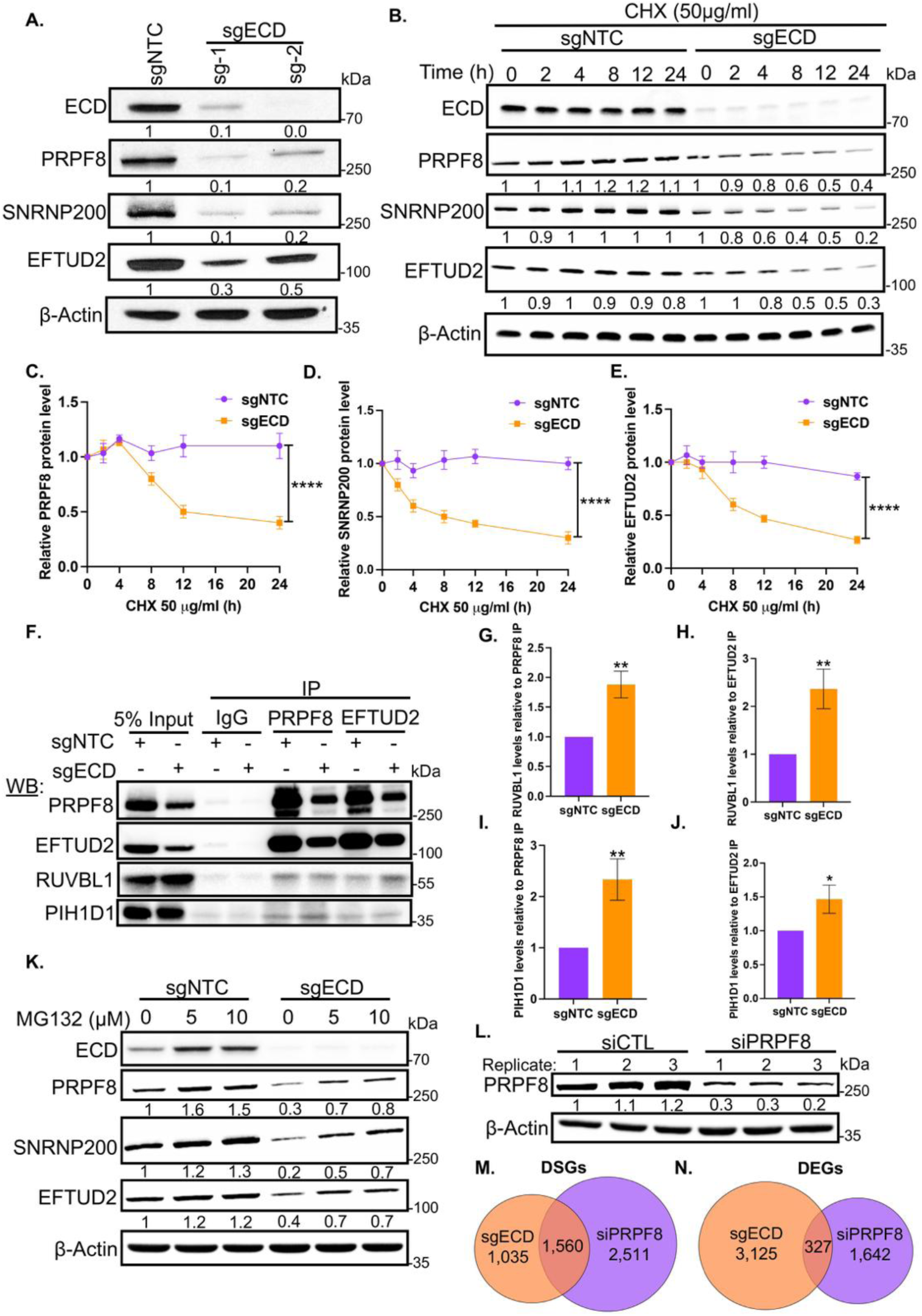
ECD depletion reduces expression of U5-specific proteins and enhances their association with the R2TP complex. **(A)** Western blot (WB) analysis of indicated protein levels in 76NTERT cells with NTC or ECD-targeted sgRNAs (sg-1/2) following doxycycline-induced Cas9 activation. **(B)** Representative western blots showing expression of indicated proteins over time following cycloheximide (CHX) treatment in control (shNTC) and ECD-depleted (sgECD) 76NTERT cells. **(C-E)** Protein decay of indicated proteins, quantified using Image J and normalized using actin and plotted over time. The data presented as mean values ± SEM from three independent experiments. **** indicate p-value <0.0001, computed through two-way Anova. **(F)** IPs followed by WB showing association of R2TP components (RUVBL1 and PIH1D1) with U5-specifc proteins (PRPF8 and EFTUD2) in control and ECD-depleted groups. **(G-J)** Quantification of RUVBL1 and PIH1D1 levels normalized to PRPF8 and EFTUD2 pull-down levels in control and ECD-depleted cells. The data presented as mean ± SEM of three independent experiments. ** p<0.01; * p-value <0.05, computed through unpaired student t test. **(K)** WB analysis of indicated proteins following mentioned dosage of MG132 treatment for 24 h. **(L)** WB showing PRPF8 levels post transfection of 76NTERT cells with control siRNA or PRPF8 siRNA. Numbers below all the blots depict band intensities in respect to loading control β-Actin, determined using Image J. **(M&N)** Venn diagrams showing overlap of **(M)** differentially spliced genes (DSGs) and **(N)** differentially expressed genes (DEGs) after depletion of ECD and PRPF8.

Notably, treatment with the proteasome inhibitor MG132 partially restored the levels of all three tested U5 proteins in ECD-depleted cells **(Fig. 7K**), suggesting that lack of ECD targets these proteins to proteosome mediated degradation. Notably, in contrast to findings reported in the *Drosophila* study (22), treatment with the lysosomal inhibitor bafilomycin A did not rescue the expression of U5 proteins in ECD depleted cells **(Fig. S6),** indicating that lysosomal degradation does not play a significant role in the mammalian cell models examined here.

Given the downregulation of U5 snRNP components upon ECD depletion, we next asked whether the observed splicing alterations are mediated through loss of U5 snRNP function or may also reflect an independent RNA-binding role of ECD. To test this, we performed siRNA-mediated knockdown of PRPF8 in 76NTERT cells **(Fig. 7L),** followed by RNA-seq analysis, and compared the DSGs and DEGs profiles with those observed upon ECD depletion. Notably, we observed a substantial overlap in DSGs between ECD-depleted and PRPF8 knockdown cells (1,560 DSGs) **(Fig. 7M)**. However, a considerable number of DSGs were unique to ECD depletion (1,035) or PRPF8 knockdown (2,511) **(Fig. 7M).** In contrast, the overlap in DEGs between ECD depleted and PRPF8 knockdown cells was limited (327 genes) **(Fig. 7N)**, with most DEGs observed exclusively in ECD depleted cells (3,125) or PRPF8 knockdown cells (1,642) **(Fig. 7N).** These findings suggest that ECD regulates RNA processing through both PRPF8-dependent and PRPF8-independent mechanisms. Furthermore, our RNA sequencing analysis revealed reduced levels of the mRNA encoded by *RNU5A-1* gene, which encodes the essential RNA component (U5 snRNA) of the U5 snRNP complex, upon ECD depletion **(Fig. S7A).** This finding was further validated through qPCR analysis of independent samples **(Fig. 8A).** We, therefore, investigated the possibility that ECD, as an RNA binding protein, may directly interact with U5 snRNA. EMSA with a biotinylated full-length U5 snRNA as the probe demonstrated strong binding of WT ECD to U5 snRNA, whereas the RNA binding mutant exhibited markedly reduced binding. Moreover, the addition of a monoclonal anti-ECD antibody effectively abrogated the ECD-U5 snRNA interaction **(Fig. 8B).** We also quantified the interaction of ECD vs. its RNA binding mutant with the U5 snRNA probe comprising 41 nucleotides from the 5′ end using the fluorescence polarization assay and observed results consistent with the EMSA data. Specifically, the WT ECD protein displayed stronger binding (Kd 3.4 ± 0.8 µM), whereas the mutant showed a marked reduction in binding (Kd 9.7 ± 2.2 µM) **(Fig. 8C).** Notably, GST pull-down assay demonstrated that the RNA binding mutant ECD retained its ability to bind to PRPF8, EFTUD2, DDX39A, RUVBL1 and RB proteins, previously identified as ECD interacting proteins, comparably to WT ECD **(Fig. 8D),** ruling out any gross structural misfolding of the ECD protein upon deletion of aa 135–148. This result also indicates that ECD can associate with protein components of the U5 snRNP complex independently of RNA. To further validate this observation *in vivo*, we performed pulldown assays in 76NTERT cells with or without RNase A treatment. We observed comparable levels of U5-specific proteins pulled down by ECD in untreated vs. RNAse-treated lysates **(Fig. S7B)**, confirming that ECD directly associates with several proteins of the U5 snRNP complex, independent of RNA.

**Fig. 8.**
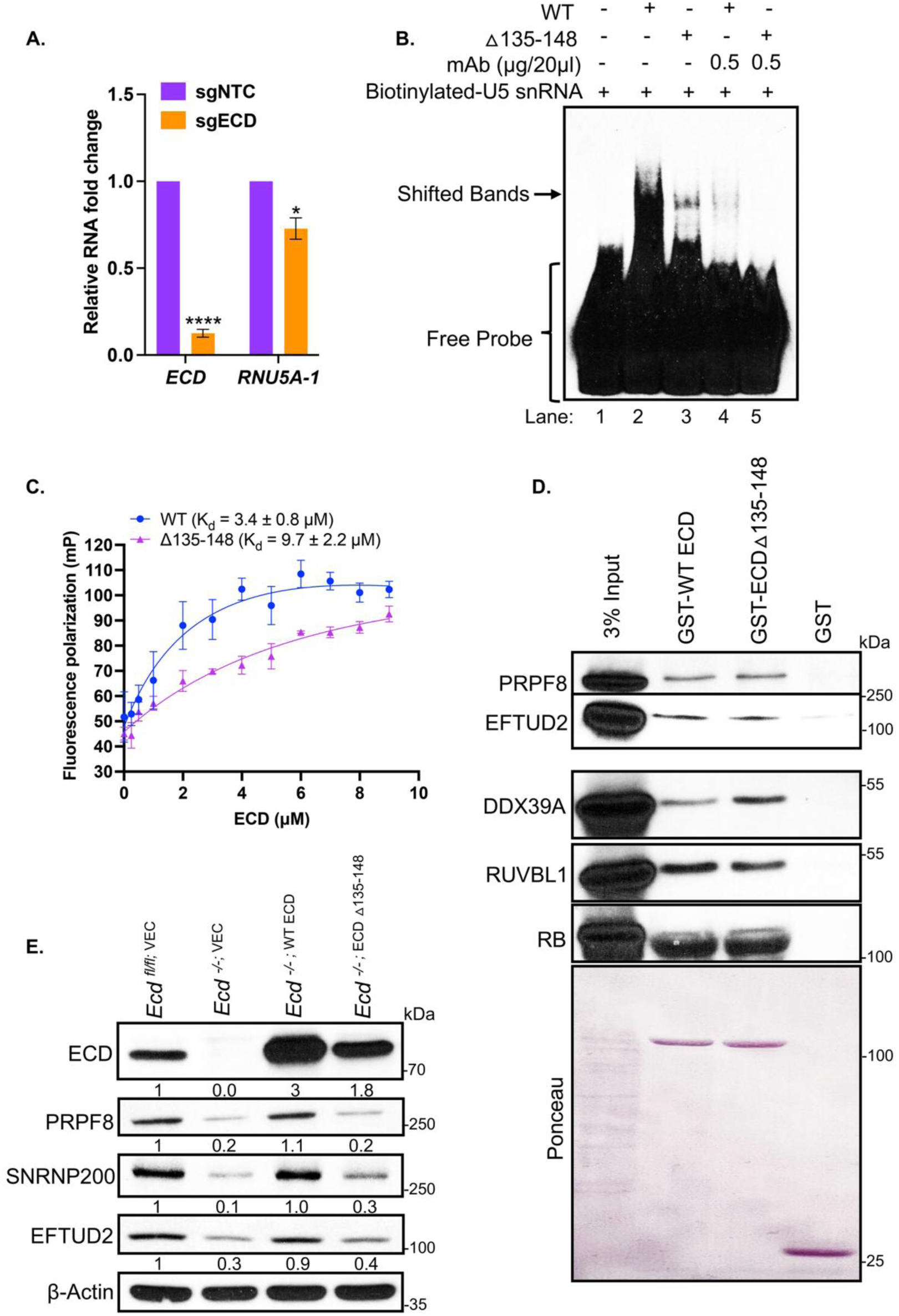
ECD binds U5 snRNA, and its RNA-binding activity is critical for U5 snRNP component expression. **(A)** Relative fold change in expression of *ECD* and *RNU5A-1* transcripts in control (sgNTC) and ECD depleted cells (sgECD), as determined through qPCR experiments.18s rRNA used as normalizing control. The data presented as mean ± SEM of three independent experiments. **** indicate p-value <0.0001; * p-value <0.05, computed through unpaired student t test. **(B)** RNA-EMSA with full length U5 snRNA as probe along with 2.5 µg/20 µl each of WT or RBD mutant of ECD in the presence or absence of ECD mAb. Experiment was performed as indicated (Fig. 1). **(C)** Fluorescence polarization assay depicting the binding affinity of WT and Δ135-148 mutant ECDs to U5 snRNA probe (5’-1-41 nucleotides) **(D)** WB shows ECD and △135-148 mutant associated proteins using GST pull down assay. 20 µg of GST-tagged WT-ECD, its RBD mutant or GST alone (negative control) immobilized on glutathione-Sepharose beads were incubated with 1 mg of protein lysate from 76NTERT and associated proteins were analyzed by WB with the indicated antibodies. The membrane was stained with Ponceau dye to visualize the quantity of fusion proteins used for pulldowns. **(E)** WB shows expression of indicated proteins in control or *Ecd* KO MEFs and upon rescue with either WT human ECD or △135-148 mutant. Numbers below the blots depict band intensities in respect to loading control β-Actin, determined using Image J.

To further explore the biological significance of ECD’s interaction with RNAs, we utilized *Ecd^fl/f^* mouse embryonic fibroblasts (MEFs) which allows endogenous *Ecd* deletion by adenovirus-mediated Cre recombinase expression (9). As expected, endogenous mouse *Ecd* KO with Cre adenovirus resulted in the downregulation of PRPF8, SNRNP200 and EFTUD2 protein levels **(Fig. 8E)**. Significantly, while full length human ECD restored the levels of PRPF8, SNRNP200 and EFTUD2 proteins, the Δ135-148 mutant was defective **(Fig. 8E)**, despite its ability to pull down the U5 snRNP complex proteins from cell lysates (**Fig. 8D).** These findings strongly support the critical role of the ECD-RNA interaction in maintaining the stability of key U5-specific proteins.

### The RNA binding activity of ECD is critical for its requirement for cell proliferation

We have shown that ECD is required for cell proliferation in diverse cellular contexts (9,11). To assess if the RNA binding activity of ECD is required for this function, we assessed the ability of WT or Δ135-148 mutant of human ECD to rescue *Ecd^fl/fl^* MEFs from proliferative block imposed by the deletion of endogenous mouse *Ecd* (9). KO of endogenous mouse *Ecd* and expression of exogenous human ECD or its mutant protein were confirmed through WB **(Fig. 9A)**. Although the level of the mutant ECD was lower than that of the WT ECD, it was about 1.4 times higher than the endogenous mouse Ecd **(Fig. 9A).** As expected, endogenous *Ecd* deletion in *Ecd^fl/fl^* MEFs resulted in a proliferation arrest, determined through measuring proliferation rate using cell-titre glo cell viability assay (**Fig. 9B)** and colony formation assay **(Fig. 9C & D).** Notably, this proliferation block was significantly rescued by WT human ECD expression (**Fig. 9B-D**). In contrast, ectopic expression of the Δ135-148 mutant failed to rescue the proliferation arrest **(Fig. 9B-D)**. These results support the conclusion that the RNA binding activity of ECD is required for its broader biological function in a cellular proliferation context.

**Fig. 9.**
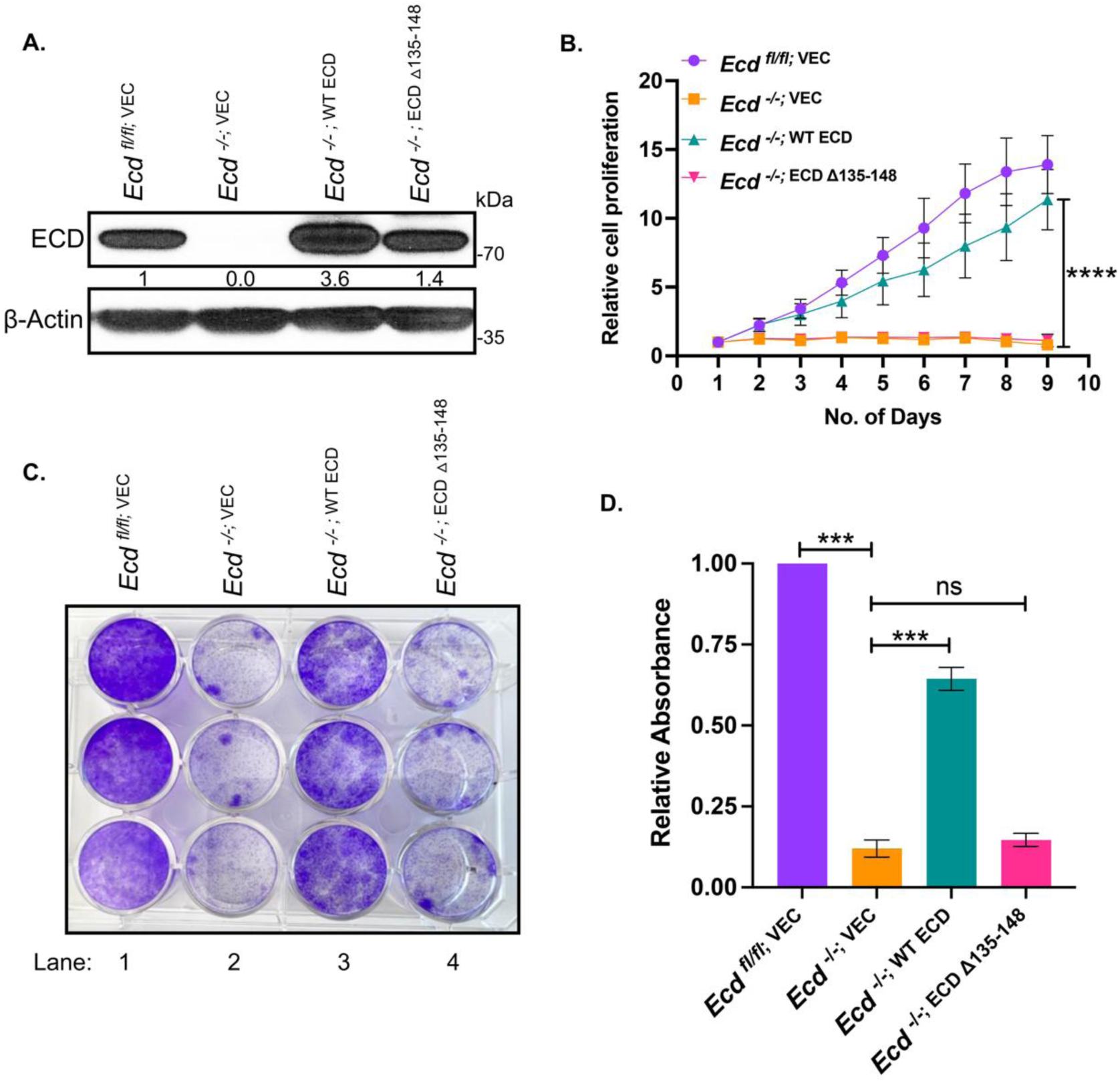
RNA binding defective mutant fails to rescue proliferation block caused by deletion of endogenous ECD. **(A)** Western blotting shows the expression of WT human ECD or its deletion mutant in *Ecd^fl/fl^* mouse embryonic fibroblasts (MEFs) with control or Cre adenovirus infection. Numbers below the blot depict band intensities in respect to loading control β-Actin, determined using Image J. **(B)** Proliferation rate of control or ECD deleted MEFs without or with rescue by WT ECD or its △135-148 mutant. *Ecd^fl/fl^* MEFs expressing vector (VEC), WT ECD or its △135-148 mutant were infected with vector or Cre adenoviruses to delete endogenous Ecd, followed by cell viability measurement at the indicated time points using CellTiter-Glo based luminescent assay. The data presented as mean values ± SEM from three independent experiments. **** indicate p-value <0.0001, computed through two-way Anova. **(C)** Representative images of colony forming abilities of *Ecd^fl/fl^* MEFs expressing full-length WT ECD or △135-148 mutant post infection with vector or Cre adenoviruses, colonies were stained with crystal violet after 10 days, and the solubilized dye absorbance was read at 590 nm. **(D)** Histograms show the relative rescue efficiency of WT ECD and △135-148 mutant as compared to vector control cells. The data represents mean ± SEM from three independent experiments. *** indicate p-value <0.001, ns= non-significant, computed through unpaired student t test.

## Discussion

ECD is an evolutionarily conserved protein with critical developmental and physiological roles and whose overexpression contributes to promoting oncogenic behaviors in multiple cancers (12–16,18). Recent studies have linked ECD to fundamental processes in RNA biogenesis, including the regulation of mRNA splicing (22). However, the intrinsic biochemical activities of ECD that underlie its roles in mRNA splicing and the linkage of such activities to its cellular roles remain unclear. Here, we identify ECD as a novel RNA binding protein that directly binds to a large subset of cellular mRNAs as well as to U5 snRNA to regulate mRNA splicing. We demonstrate that ECD’s RNA binding activity requires the structurally conserved N-terminal region. RNA binding defective mutant (△135-148) lacks the ability to stabilize the three tested proteins of U5 snRNP complex or to rescue the cell proliferation. Further, we provide evidence that association of ECD to protein components of U5 snRNP complex is RNA independent. Our findings provide novel mechanistic insights into an essential cellular regulator of mRNA splicing whose biochemical roles have been nebulous thus far.

We provide multiple lines of evidence supporting ECD’s direct RNA binding function. These include in silico RNABindRplus prediction of potential RNA binding motifs on ECD and validation of RNA binding by EMSA and FP analyses **(Fig. 1A-F)**. Selective deletion of aa 135-148 in ECD significantly reduced its RNA binding capacity **(Fig. 1E-F)** but did not alter its interactions with proteins known to be associated with other regions of ECD, including PRPF8, EFTUD2, DDX39A, RUVBL1 and RB **(Fig. 8D).**

While ECD’s structure has not been determined experimentally, our circular dichroism (CD) and small-angle X-ray scattering (SAXS) studies (23) and AlphaFold modeling (25) support a well-folded structure of the N-terminal region of ECD where the RNA binding critical amino acids are located **(Fig. 1G)**, whereas the C-terminus is disordered, containing intrinsically disordered regions (IDRs). Notably, IDRs are a characteristic of many RBPs where they confer structural flexibility to enhance the RNA interaction potential (49). RNA chaperones often utilize IDRs as functional domains for dynamic RNA interactions and to facilitate RNA folding or processing (50,51). While our analyses show that IDR containing C-terminal regions of ECD are dispensable for RNA binding, it remains possible that the IDR containing C-terminal regions may facilitate RNA binding *in vivo*, potentially by enhancing ECD’s conformational flexibility or interaction dynamics within the cellular environment or through protein-protein interactions. In this context, it is notable that the previously defined interactions of ECD with proteins like PIH1D1 component of the R2TP co-chaperone complex and with the cell cycle regulator RB are mediated by C-terminal regions (9,10). Significantly, while the *Drosophila* Ecd sequence (aa 160–173) corresponding to human ECD aa 135–148, which we identify as important for RNA binding, does not exhibit high conservation at individual amino acids but are predicted by the AlphaFold to form identical anti-parallel beta sheet structures in both mammalian and fly ECD proteins **(Fig. S2).** Given the known importance of anti-parallel beta sheets in RNA recognition motifs (RRM) of RNA binding proteins (52), it is likely that the region we identify as critical for RNA binding activity of human ECD will encode a conserved RNA binding activity across species.

Our eCLIP-seq analyses provided evidence for ECD’s broad role as an RBP, demonstrating 3,363 unique transcriptome-wide binding sites on mRNAs encoded by genes involved in essential cellular processes, including transcription, apoptosis, endoplasmic reticulum stress, cellular differentiation, and proliferation **(Fig. S3A).** These results are consistent with roles of ECD in physiological processes such as cell proliferation (9) and cell survival under endoplasmic reticulum stress (11) as well as the role of ECD overexpression in promoting tumorigenesis (13–16,19).

Notably, mapping the RNA binding preferences of ECD revealed a strong preference for coding sequences (CDS) and 5’ untranslated regions (5’UTRs) of target RNAs. The top ECD binding motif present on a fraction of target RNAs identified as a purine rich sequence, GAGGAGGAGGAG, that includes the sequence of GAGGA, previously reported to present in exonic splicing enhancers (ESE) (53). ESE sequences are typically purine-rich and often consist of motifs alternating between adenosine (A) and guanosine (G), spanning six or more nucleotides. Their purine-rich composition is known to enhance the binding of splicing factors and facilitate RNA splicing (54). However, in vitro, we observed slightly weaker binding to this sequence as compared to the pyrimidine rich sequence **(Fig. 2H&I)**, suggesting that ECD–RNA binding selectivity in vivo may be further influenced by associated proteins, especially since several ECD associated proteins (e.g., the U5 snRNP components) have an ability to directly bind to RNAs (19). Consistent with the functional importance of ECD binding to splicing relevant motifs in target RNAs in cellular context, depletion of ECD resulted in thousands of differential splicing events, especially causing alteration in exon skipping and intron retention features **(Fig. 4A-C).** Significantly, exonic ESE sequences within viral and mammalian genes are known to play a critical role in regulating alternative RNA splicing (55). Notably, we previously found ECD to be important in regulating viral and cellular RNA splicing in the Human papillomavirus (HPV) positive, cervical cancer cell line model (13).

While we observed an enrichment of the ECD binding peaks in RNAs proximal to mis-spliced junctions **(Fig. 5C),** a diffuse distribution of ECD binding across the length of transcripts was also seen **(Fig. 5D &E, and Fig. 6D)**. This pattern is consistent with RNA-binding proteins that exhibit flexible or low-sequence-specific binding behavior **(Fig.6D)** (56). These findings suggest that ECD association with transcripts does not rely exclusively on strict sequence recognition but is likely influenced by context-dependent protein interactions. Such interactions may modulate splicing outcomes through cooperation with other splicing factors. However, to pinpoint the detailed mechanism by which ECD regulates splicing through direct RNA binding will require extensive further investigation. Notably, only about 22% of the differentially spliced RNAs are directly bound to ECD, suggesting that ECD regulates splicing through additional mechanisms beyond direct binding to splicing sites or regulatory elements. One of these mechanisms is likely to be the requirement of ECD for proper splicing of transcription factors and accessory proteins that regulate transcription, splicing, stability, and other aspects of RNA biogenesis. In this regard, a significant subset of genes whose transcripts showed aberrant splicing were transcription factors whose target gene mRNA levels were altered (**Fig. 5F & G**). This explains one of the mechanisms through which ECD regulates the expression of thousands of genes, including both coding and non-coding RNAs **(Fig. 3C & S3C).** However, it also impacts gene expression for some of its targets through direct RNA binding, as transcripts of 6% of total DEGs were identified to bind directly to ECD. Our observation that the ECD-bound transcriptomic profile overlaps with the eCLIP-seq profiles of PRPF8 and EFTUD2 (**Fig. 6A-D**), two protein components of the U5 snRNP complex that associate with ECD suggests that the RNA binding specificity of ECD *in vivo* is not strictly RNA sequence-driven and potentially context dependent, influenced by protein interactome. The absence of strict RNA sequence specificity of ECD aligns with certain previously reported RBPs involved in splicing regulation, which exhibit minimal or no sequence selectivity, recognizing RNA structures or being influenced by factors such as protein cofactors (37,57).

We and others have previously reported that ECD engages in protein-protein interactions with components of the RNA biogenesis and translation machinery, with several proteins of the U5 snRNP complex in the ECD-associated proteome (19,22). Such protein-protein interactions have suggested one plausible mechanism for ECD to play a role in RNA splicing, with recent evidence that *Drosophila* Ecd functions as a chaperone to stabilize the U5 snRNP during its cytoplasmic assembly prior to nuclear import (22). Our results show that, like *Drosophila* Ecd (22), the mammalian ECD protein is required to help maintain the stability of the U5 snRNP complex protein PRPF8, and we extend this to other U5-specific proteins including the SNRNP200, an RNA helicase, and EFTUD2, a GTPase **(Fig. 7A-E).** Both proteins help dissociate the U4 and U6 snRNPs during splicing to facilitate the processing of precursor mRNAs into mature mRNAs (58,59). Mechanistically, upon ECD depletion, we observed increased association of these proteins with R2TP components RUVBL1 and PIH1D1 **(Fig. 7F-J & S5A-E)**, R2TP components known to associate with ECD (10) and reported to be involved in U5 snRNP assembly (60,61), suggesting that ECD and its interaction with the R2TP complex play a critical role in the early assembly and stabilization of the U5 snRNP. The enhanced interaction of U5 snRNP proteins with the R2TP complex in ECD-depleted cells likely reflects the requirement of ECD in a post-R2TP assembly step, with the loss of ECD resulting in the R2TP cochaperone targeting of unassembled or partially assembled U5 components for proteasome-mediated degradation **(Fig. 7K)**. Supporting this model, PRPF8 mutants that fail to properly integrate into U5 snRNP also exhibit stronger binding to R2TP (61), indicating that R2TP may preferentially engage the unassembled U5 components. Together, these findings suggest that ECD is necessary for efficient U5 snRNP assembly and for preventing prolonged retention of its components by cytoplasmic chaperones, thereby ensuring proper maturation and splicing function. In addition, ECD may also help recruit key additional proteins to help assemble and/or stabilize the U5 snRNP complex. For example, ECD was shown to assemble into a complex that included PRPF8, R2TP and additional proteins including AAR2 (62). AAR2 itself is a critical factor in U5 snRNP assembly (63). Furthermore, we show that ECD directly interacts with U5 snRNA **(Fig. 8B&C),** the essential RNA component of the U5 snRNP complex (64). Importantly, while the WT ECD rescued the protein expression of PRPF8, SNRNP200 and EFTUD2 observed upon ECD loss, the RNA binding mutant ECD failed to do so **(Fig. 8E)**, thus establishing the RNA binding activity of ECD as pivotal for maintaining the expression of these U5 proteins. It is notable that the prior work showed the *Drosophila* Ecd protein to co-IP with U5 snRNA (22) but a direct interaction of fly Ecd with the SmD3 component of the U5 Sm Ring implied that co-IP of Ecd and U5 snRNA may be indirect. Our results demonstrate the direct ECD binding to U5 snRNA and regulation of key proteins of U5 snRNP complex (PRPF8, SNRNP200 and EFTUD2), supporting a model in which ECD stabilizes a fully assembled and functional U5 snRNP complex. ECD’s RNA-independent association with the U5 snRNP protein components **(Fig. S7B)**, concurrent binding to its RNA component, U5 snRNA, and its role in maintaining the U5 proteins stability, suggests that ECD functions as a central regulator of U5 biogenesis and splicing, potentially acting as a molecular scaffold to promote assembly and stability. Consistent with this model, mutations or loss of U5 snRNA in yeast significantly reduce the Prp8 protein levels emphasizing the interdependence between snRNAs and spliceosome proteins (65). Pertinent to ECD’s dual interaction with RNA and protein components of complex molecular machines like the U5 snRNP, it will be of great interest to explore if ECD plays a role in the assembly of other RNA-protein complexes, such as the small nucleolar ribonucleoprotein (snoRNP) complexes, which are well-established to require the ECD interacting R2TP complex for their assembly (66).

While our studies are consistent with the role of ECD in the cytoplasmic assembly of U5 snRNP, as shown for *Drosophila* Ecd (22), the direct binding of ECD to a large repertoire of mRNAs in exonic regions near their splice junctions supports its role in splicing beyond the U5 snRNP assembly. In this regard, knockdown of PRPF8 in the same cellular system followed by RNA-seq analysis **(Fig. 7L)** revealed both shared and distinct DSGs and DEGs with ECD depletion **(Fig. 7M&N)**. Notably, a substantial number of DSGs or DEGs were specific to either ECD or PRPF8 depletion, suggesting that ECD may also exert PRPF8-independent roles in RNA processing. While ECD regulates the stability of PRPF8 and other U5-specific proteins, these results suggest that ECD function in splicing likely extends beyond the early U5 snRNP assembly. Importantly, we have found ECD to shuttle between the cytoplasm and nucleus (67). Whether ECD shuttling might represent its necessity to maintain an intact U5 snRNP complex even during the steps of RNA splicing in the nucleus, is an important question that needs further exploration.

Functional assay with cell proliferation as a readout demonstrate that the RNA binding deficient mutant of ECD failed to rescue MEF cells from a proliferative block imposed by the endogenous Ecd KO **(Fig. 9B-D),** highlighting the importance of RNA binding for its cellular function. Collectively, our findings elucidate novel aspects of the mechanistic basis for ECD’s role in maintaining the fidelity and efficiency of pre-mRNA splicing. Given the critical physiological functions of ECD and the pro-oncogenic effects due to its overexpression in cancers, these new mechanistic insights add a novel perspective in understanding its roles in cellular and cancer biology and devising means to disable ECD’s function for therapeutic purposes.

## Data availability

The e-CLIP and RNA sequencing data generated in this study are deposited under Gene Expression Omnibus (GEO) accession: GSE285409.

## Acknowledgements

This research was funded by Pilot grants from the Fred & Pamela Buffett Cancer Center (HB & VB); Department of Defense grants W81XWH-17-1-0616 and W81XWH-20-1-0058 to HB and W81XWH-20-1-0546; HT94252410337; and National Institutes of Health (NIH) grants R21CA241055 and R03CA253193 to VB; the NIH MIRA award R35 GM147467 (to MJR); the Raphael Bonita Memorial Fund and support to UNMC core facilities from the NCI Cancer Center Support Grant (P30CA036727) awarded to Fred & Pamela Buffett Cancer Center and from the Nebraska Research Initiative.

## Author contributions

Conceptualization: MR, HB, MJR and VB. Methodology, Data Analysis and Interpretation: MR, ARR, AK, BM, IS, BBK, KKB, HB, MJR, and VB. Writing – original draft: MR. Writing – review and editing: MR, ARR, AK, BM, IS, BBK, KKB, HB, MJR, and VB. Bioinformatic analyses: MR, AK and MJR. Supervision: HB, MJR, and VB. Funding acquisition: HB, MJR, and VB.

## Supplementary Data Statement

Supplementary Data are available at NAR online.

