## Supplemental File for "A novel RNA-binding activity of ECD contributes to U5 snRNP stability and pre-mRNA splicing"

**Supplemental Figure S1**

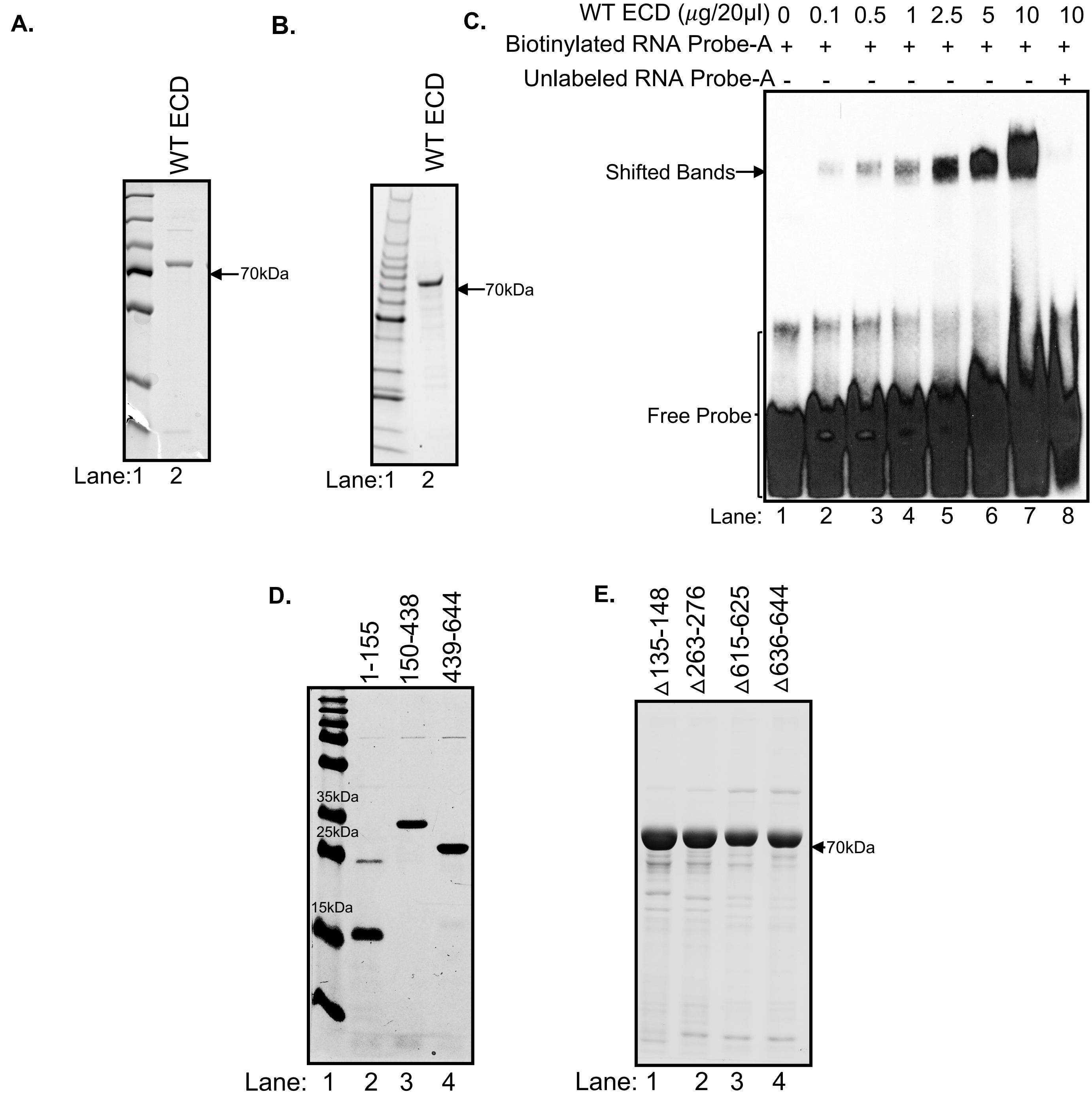

**Supplemental Fig. S1. Assessment of purity of recombinant WT ECD and various mutants and RNA binding capacity of His-tagged ECD.** Coomassie stained SDS-PAGE gels show the purity of **(A)** GST affinity purified and **(B)** Ni-NTA affinity purified His tagged wild type (WT) ECD proteins. **(C)** RNA-EMSA with different concentrations of purified His tagged ECD (lanes 2 to 7) along with biotinylated RNA probe-A (40 nM). Protein free reaction mix, used as a negative control (lane 1). Competition with 100-fold excess unlabeled probe was done to determine the binding specificity (lane 8). **(D)** SDS-PAGE gels showing the purity of GST affinity purified truncated mutants and **(E)** indicated deletion mutants of ECD.

### Supplemental Figure S2

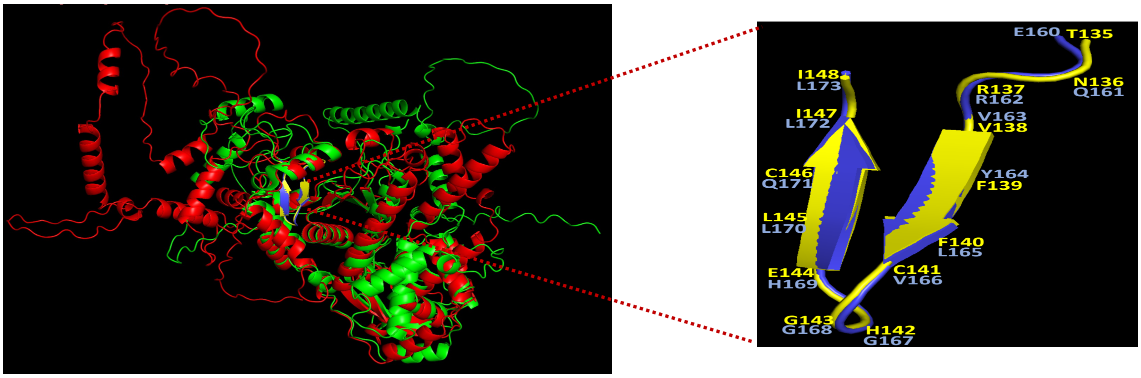

**Supplemental Fig. S2. Corresponding amino acids in *Drosophila* Ecd exhibit identical secondary structure to RNA binding region of human ECD.** Superimposed predicted tertiary structures of human ECD (green) and *Drosophila* Ecd (red) proteins, showing identified RNA binding region (aa 135-148) of human ECD protein (in yellow) with respective residues (aa 160-173) in *Drosophila* Ecd (in blue). PDB files of Alpha fold predicted 3D structures of human ECD and *Drosophila* Ecd proteins were visualized and superimposed using PyMOL software.

### Supplemental Figure S3

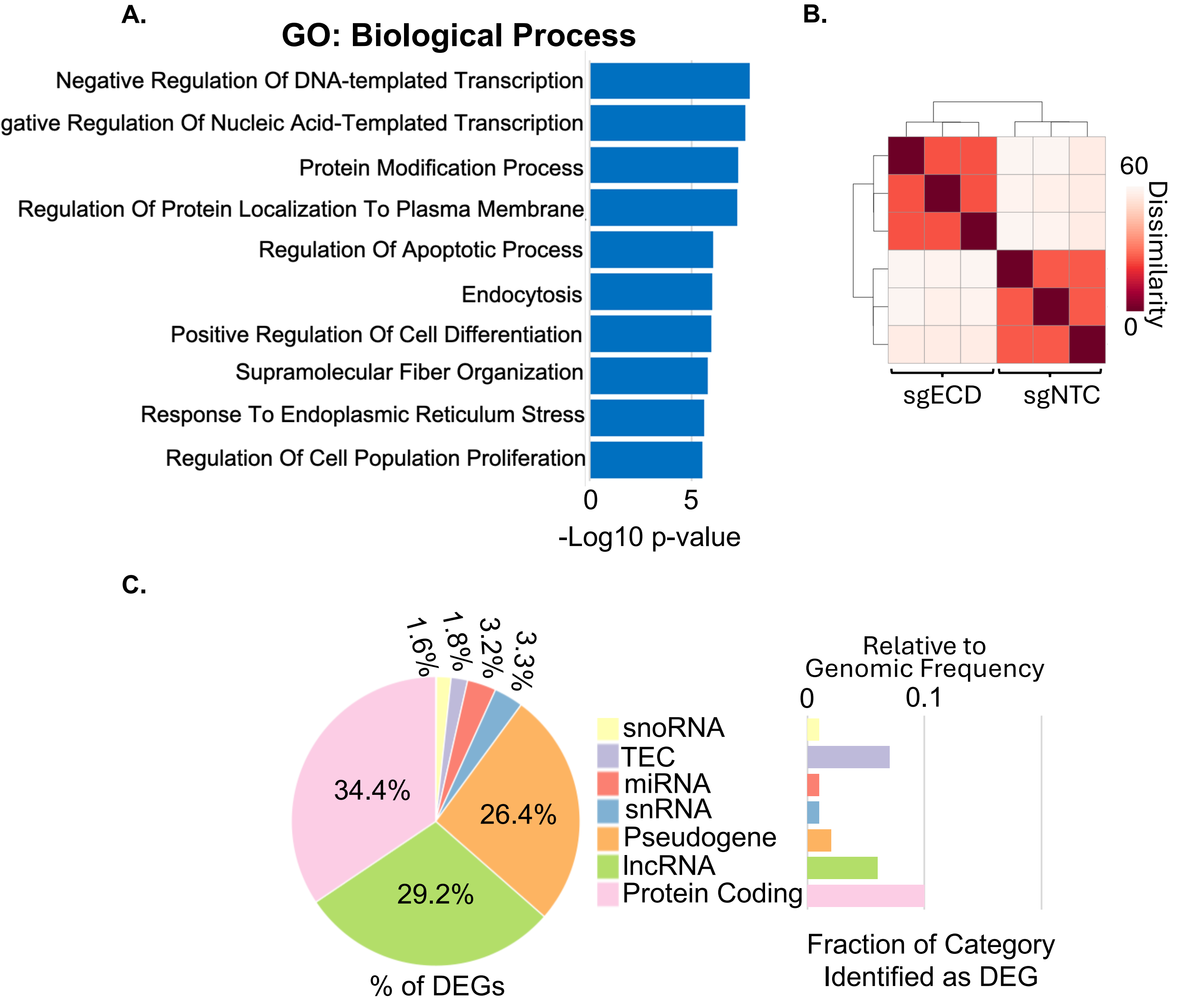

**Supplemental Fig. S3. Biological process affected by RNAs bound by ECD and RNA biotypes affected by ECD depletion. A)** Top pathways affected by genes harboring ECD binding consensus motif as identified through GO Biological pathway analysis. **(B)** Replicate comparison of RNA-seq data. **(C)** Categories of annotations for differentially expressed genes showing the raw percentages (left) and the frequency relative to their total presence throughout the genome (right).

**Supplemental Figure S4**

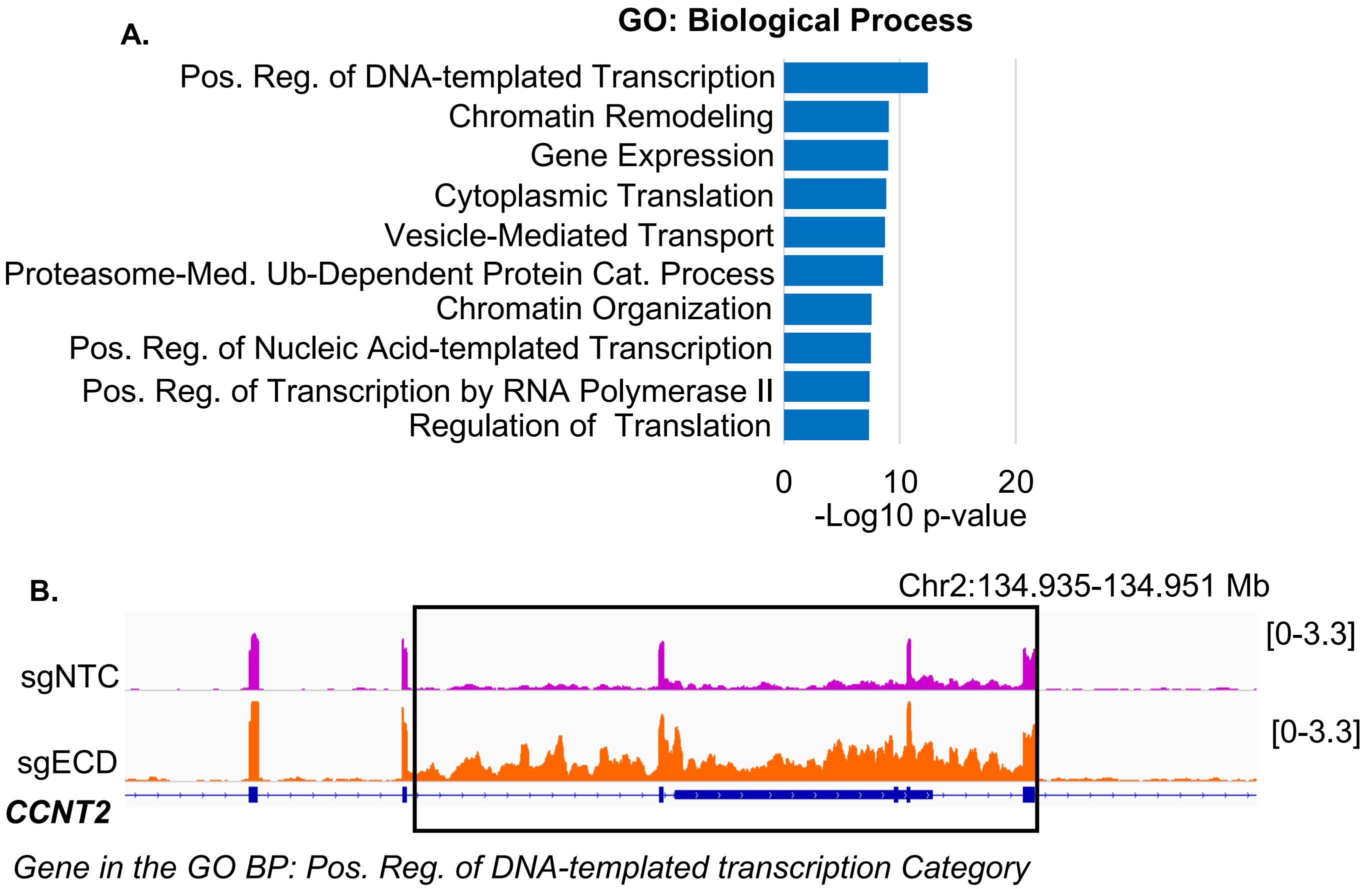

**Supplemental Fig. S4. Pathways affected by differential spliced genes caused by ECD depletion. (A)** The top gene ontology biological processes gene sets associated with differentially spliced genes after ECD depletion. **(B)** RNA-seq signal track showing an aberrant retention in transcript of a gene in the top category (shown in A) after ECD depletion.

**Supplemental Figure S5**

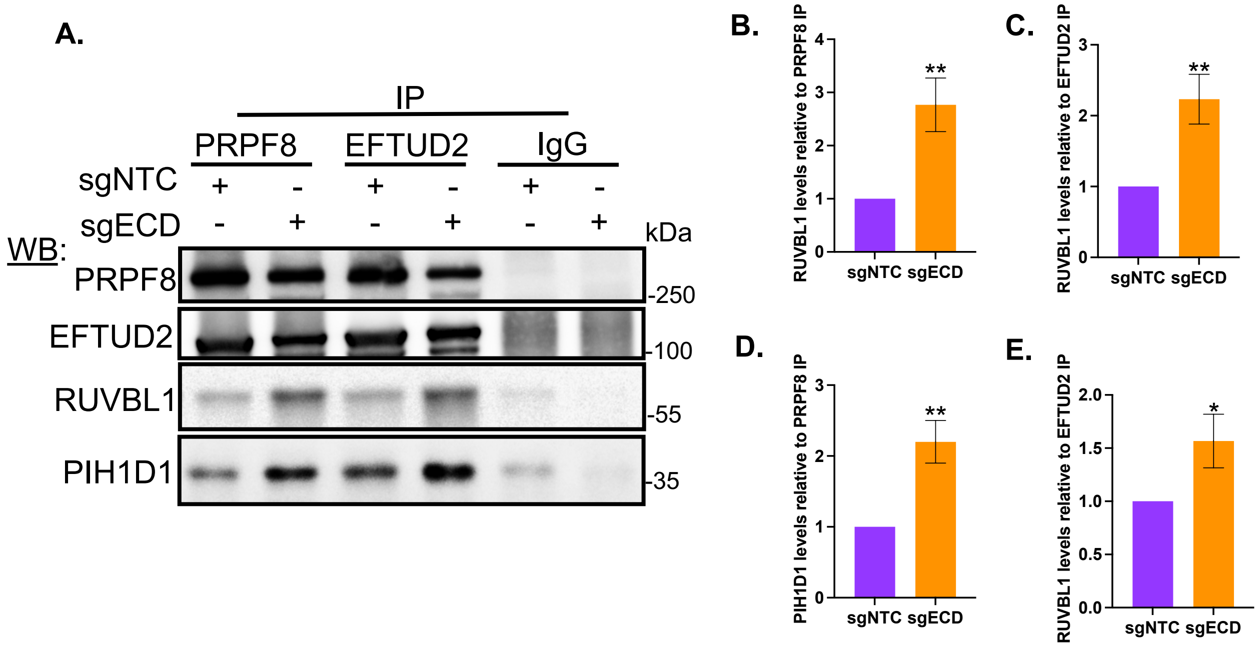

**Supplemental Fig. S5. Loss of ECD enhances U5-specific proteins interaction with R2TP components. (A)** IPs followed by WB showing association of RUVBL1 and PIH1D1 with PRPF8 and EFTUD2 in control and ECD-depleted groups. **(B-E)** Quantification of RUVBL1 and PIH1D1 levels normalized to PRPF8 and EFTUD2 pull-down levels in control and ECD-depleted cells. The data presented as mean ± SEM of three independent experiments. ** p<.01 ; * p-value <0.05, computed through unpaired student t test.

**Supplemental Figure S6**

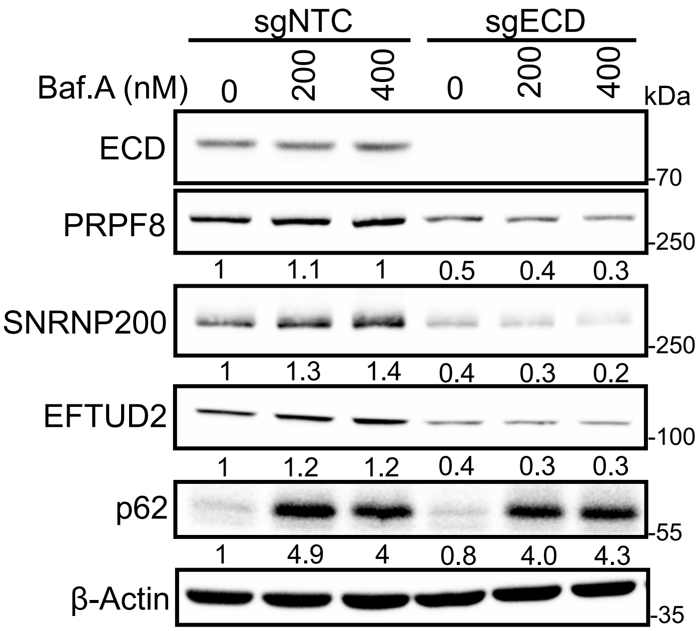

**Supplemental Fig. S6. Treatment with bafilomycin A does not rescue U5-specifc protein levels following ECD depletion.** WB analysis of U5 proteins following treatment with indicated doses of bafilomycin A (Baf.A) for 24 h. p62 was used as a positive control. Numbers below the blots depict band intensities in respect to loading control β-Actin, determined using Image J.

Supplemental Figure S7

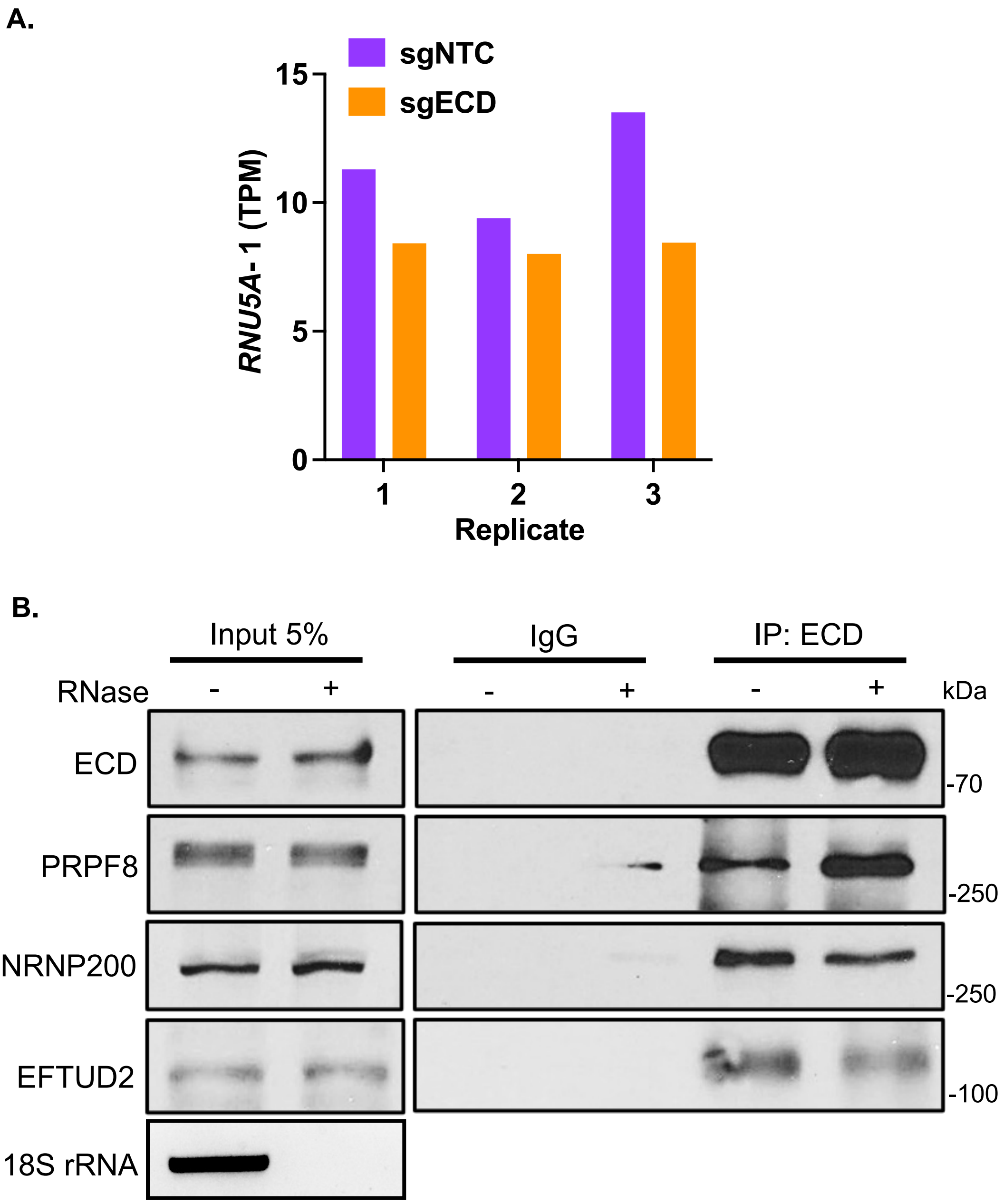

**Supplemental Fig. S7. Loss of ECD results in downregulation of U5 snRNA and ECD associates to protein components of U5 snRNP complex independent of RNA.** (A) Histograms showing expression in TPM of *RNU5A-1* transcript in three biological replicates from control and ECD depleted cells as identified through RNA seq analysis. **(B)** Representative WB of the indicated proteins after immunoprecipitation using either IgG or ECD specific antibodies in untreated or RNase A treated protein lysates from 76NTERT mammary epithelial cells. Pulldowns through ECD show comparable levels of U5 snRNP protein components in both untreated and RNase A treated conditions. Semi-quantitative PCR of 18S rRNA was used to confirm the efficacy of RNase A treatment.

**Supplementary Table S1**: Nucleotide sequence of Primers and RNA- Probes for EMSA and FP assays.

| | | | |
|---|---|---|---|
| **qPCR** | **Gene** | **Forward Sequence (5'-3')** | **Reverse Sequence (5'-3')** |
| | *ECD* | ACTTTGAAACACACGAACCTGGCG | TGATGCAGGTGTGTGCTAGTTCCT |
| | *18S* | GCTTAATTTGACTCAACACGGGA | AGCTATCAATCTGTCAATCCTGTC |
| | *RNU5A-1* | TGGTTTCTCTTCAGATCGCATA | CAAAGCAAGGCCTCAAAAA |
| **Site-directed mutagenesis** | ECDΔ135-148 | CCTGCACCAAGAAAATCTG | GCTATTTTCAGGATCCAG |
| **EMSA Probes** | **Probe Name** | **Modification** | **Sequence (5'-3')** |
| | Probe A | 5'-Biotin | AGCGUGCCGUGCAACAACAUUACAAUUUACAAUCCACCAUGG |
| | Probe B | 5'-Biotin | UCAACGACUCAACGAC |
| | U5 snRNA | 5'-Biotin | AUACUCUGGUUUCUCUUCAGAUCGCAUAAAUCUUUCGCCUUUUACUAAAGUUUCCGUGGAGAGGAACAACUCUGAGUCUUAACCCAAUUUUUUGAGGCCUUGCUUUGGCAAGGCUA |
| **FP Assay** | **Probe Name** | **Modification** | **Sequence (5'-3')** |
| | Probe A | 5'-6-FAM-Sp9 | AGCGUGCCGUGCAACAACAUUACAAUUUACAAUCCACCAUGG |
| | Purine Exclusive | 5'-6-FAM-Sp9 | GAGGAGGAGGAGGAG |
| | Purine-Pyrimidine Mix (Probe B) | 5'-6-FAM-Sp9 | UCAACGACUCAACGAC |
| | U5 snRNA_5'-41nt | 5'-6-FAM-Sp9 | AUACUCUGGUUUCUCUUCAGAUCGCAUAAAUCUUUCGCCUU |
